# Composing trajectories for rapid inference of navigational goals

**DOI:** 10.1101/2025.09.24.678123

**Authors:** Nada Y. AbdelRahman, Wanchen Jiang, Luke T. Coddington, Sheng Gong, Joshua T. Dudman, Ann M. Hermundstad

## Abstract

Animals efficiently learn to navigate their environment. In the laboratory, naive mice explore their environment via highly structured trajectories and can learn to localize new spatial targets in as few as a handful of trials. It is unclear how such efficient learning is possible, since existing computational models of spatial navigation require far more experience to achieve comparable performance and do not attempt to explain the evolving structure of animal behavior during learning. To inform a new algorithm for rapid learning of navigational goals, we took inspiration from the reliable structure of behavior as mice learned to intercept hidden spatial targets. We designed agents that generate behavioral trajectories by controlling the speed and angular velocity of smooth path segments between anchor points. To rapidly learn good anchors, we use Bayesian inference on the history of rewarded and unrewarded trajectories to infer the probability that an anchor will be successful, and active sampling to trim hypothesized anchors. Agents learn within tens of trials to generate compact trajectories that intercept a target, capturing the evolution of behavioral structure and matching the upper limits of learning efficiency observed in mice. We further show that this algorithm can explain how mice avoid obstacles and rapidly adapt to target switches. Finally, we show that this framework naturally encompasses both egocentric and allocentric strategies for navigation.

## Introduction

A hallmark of animal cognition is the ability to make efficient use of experience to navigate through the environment, adapt motor skills, and infer goals^1–4^. When foraging for food, for example, mice traverse large and complex territories to novel or known food sources before returning to their burrow. In the laboratory, mice quickly learn to navigate to hidden targets^5,6^ or around obstacles^7^ based on a handful of successful experiences. Current computational models have been remarkably successful in accounting for brain representations thought to be critical for such navigation, especially when there are sensory cues that can be used to localize oneself in space^8–13^. However, animals can deploy memory-guided navigation even in the absence of strong localizing sensory cues^4^. Moreover, current models do not (attempt to) explain the structured evolution of navigational trajectories observed during learning, and they typically require orders of magnitude more experience to build representations that enable efficient navigation and localization of goals^11,13–15^, even when provided with highly structured state spaces^16,17^. While further augmentation and refinement of current learning algorithms is reducing this gap^5,14,15^, it is also clear that animal navigation at more naturalistic scales will present even more significant challenges to the efficiency of these algorithms^18,19^. The discrepancies in the structure and efficiency of learning between models and animals suggest that new algorithmic approaches may be critical.

One place to look for algorithmic inspiration is in the structure of behavioral trajectories that animals use to navigate new settings. When mice explore a new environment, their exploration is highly structured and does not resemble random exploration that is common to many existing algorithms^5^. In particular, trajectories of freely running animals appear to be composed of two or more smooth path segments characterized by bell shaped velocity profiles^5^ joined at ‘anchor points’ or ‘subgoals’ ^7^. When changes are made to the structure of a known environment, mouse behavior also exhibits structured changes in trajectories to incorporate new subgoals^7,14^. How might learning algorithms exploit the structure of trajectories to rapidly infer and refine navigational goals?

To address this, we analyzed the behavior and learning efficiency of naive mice in a hidden-target, delayed-reward navigation task. We show that the fastest mice can learn to intercept targets within a handful of reinforcements, and they do so by rapidly refining extended multi-segment trajectories into compact ones. We then describe an algorithm that learns to modify the parameters of a motor controller based on the outcomes of past actions, and we show how this algorithm captures the evolution of behavioral structure and the upper limits of learning efficiency observed in mice. We use this conceptualization to provide algorithmic insight in several other domains. For example, when obstacles are introduced into the environment, we show that mice and agents augment their trajectories with additional stable anchor points to form new, multi-component trajectories that avoid obstacles while still intercepting the stable target. When targets are moved within the environment, we show that both mice and agents reconfigure their trajectories to intercept new targets while still making occasional directed runs that would have intercepted previously-rewarded targets. We conclude by showing how this algorithm naturally supports learning in allocentric coordinate frames.

## Results

### Mice use structured behavioral trajectories to explore space and rapidly intercept hidden targets

We first consider a behavioral task that requires mice to learn to generate navigational trajectories using only delayed feedback delivered via a home reward port in a fixed location^5^. As described previously, mice are placed in a darkened, featureless arena (~75 cm per side) with a reward port located on the south wall. In order to trigger a drop of water reward at the port, mice must briefly occupy an unmarked (hidden) target positioned tens of centimeters away from the port. Mice are free to explore the arena while returning to the reward port to assess whether their navigational trajectory successfully led to the presence of a reward at the port (schematized in Fig. 1A). Mice do not receive localizing feedback when they intercept the target, and they must instead use delayed feedback and are required to return to the vicinity of the home port to re-activate the target location (*i.e*., multiple target interceptions do not produce multiple rewards without collection; see Methods for additional details).

**Figure 1:**
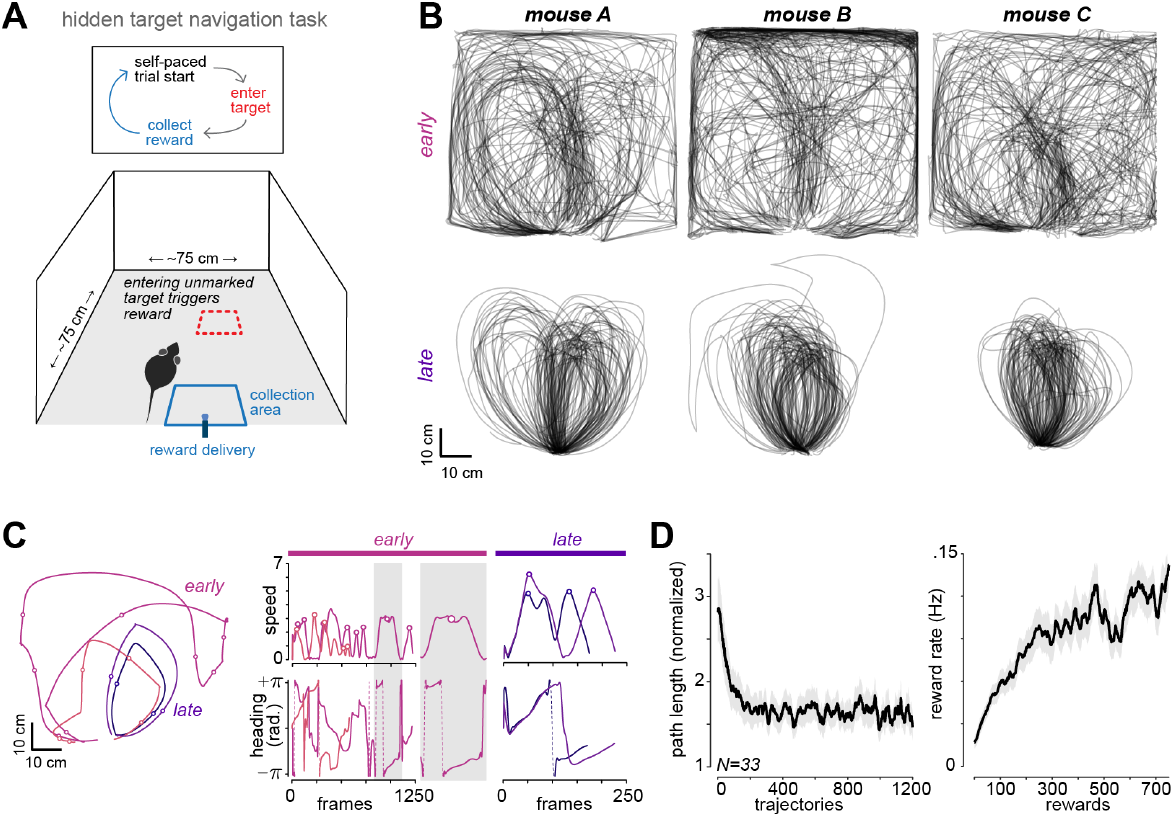
Mice rapidly refine navigational trajectories to intercept hidden targets. **A)** Schematic of the hidden target behavioral paradigm for mice. **B)** Example trajectories (100 trajectories that intercept the target) from three mice taken from the first session (“early”, naive animals, upper) and at the beginning of the seventh session (“late”, lower). **C)** To illustrate the trajectory structure, the left panel shows two example trajectories each from early and late sessions, taken from mouse C. The right panels show the instantaneous speed and heading angle along each path (note the different temporal scaling). The gray inset shows a zoomed-in portion of an early trajectory. **D)** Path length (normalized to edge length of arena) and reward rate for a population of 33 mice exhibiting consistent, rapid learning.

Populations of mice exhibit substantial variance across individuals, but at the upper limit (in a cohort of >30 individuals), mice can adapt to novel target locations with only tens of reinforced actions (Fig. 1B). All mice naive to the task exhibit consistent changes in behavior over the first ~100 reinforcements. Figure 1B shows trajectories that intercepted the target for three example mice. During the first session of the task (<100 reinforcements), their trajectories are complex and cover the arena approximately uniformly (Fig. 1B, “early”). By the beginning of the seventh session, these same mice, like all other mice tested, produce stereotyped trajectories that reliably intercept the target via smooth and efficient loops (Fig. 1B, “late”). To further illustrate these changes in trajectory structure, we can examine a pair of neighboring trajectories from early versus late epochs (Fig. 1C). Both early and late trajectories are characterized by peaks in running speed that can be used to partition trajectories into segments; these peaks in running speed are accompanied by approximately linear changes in heading. Early trajectories are long (Fig. 1C-D, left), with many segments that vary in speed and heading (Fig. 1C, middle). As mice become more adept at triggering reward by running trajectories that intercept the target (Fig. 1D, right), their trajectories become shorter (Fig. 1C-D, left) and consist of a smooth outward and return segment (Fig. 1C, right).

This learning from tens to hundreds of rewards is several orders of magnitude faster than standard reinforcement learning models applied to trajectory learning problems^5,14^. For example, in an extensive survey of canonical reinforcement learning models applied to navigational trajectories in a similar setting although without a hidden target, Shamash et al. found that it required 35,000 (Tabular Q-learning), 45,000 (Tabular SARSA), and 125,000 (Successor representation) reinforced epochs for models to learn efficient trajectories that mice learn within tens of trials^14^. Moreover, the evolving structure of trajectories observed in such models lacks the clear, reliable structure apparent in mouse trajectories^5^. This raises the question of how it is possible for mice to learn reliable target interception strategies in the absence of clear landmarks, with delayed rewards, and with so few and so variable rewarded trajectories. While innovations such as “practice runs”—in which the experimenter shows the agent the correct path^14^—substantially improved training efficiency, agents still required more than 2,000 reinforced epochs to learn efficient trajectories^14^. Thus, even when provided the correct answer, canonical algorithms are still more than an order of magnitude less efficient than animals. We sought to develop new algorithmic approaches to address some potentially fundamental limitations in canonical reinforcement learning models of navigation and capture the evolving structure of behavior over learning.

### A minimal agent captures the evolving structure of mouse behavior during rapid learning

To understand the algorithmic basis of this rapid learning in animals, we did not seek to capture the diverse sources of variance that can be observed within individual mice (*e.g*. task disengagement^20,21^ or putative motor noise^5,22^). Rather, we sought to construct an optimal agent that captures the evolving structure of behavior while simultaneously achieving the upper limit of learning efficiency. In constructing this model, we chose not to adopt the tabular representation of behavior that is common to existing models (SI Fig. 1A). Instead, we constructed a parameterized generative model of navigational trajectories that is informed by the structure of mouse behavior, and we formulated learning in the space of trajectory parameters (SI Fig. 1B). To this end, we generate multi-segment trajectories that are defined by a set of spatial ‘anchor points’ (Fig. 2A, upper row). The vector between a pair of anchor points can then be used to derive the parameters of a motor controller that generates smooth path segments between them (Fig. 2A, middle row). When coupled together, a set of vectors between anchor points, together with an initial heading direction, fully specifies the curvilinear trajectory that smoothly and efficiently links anchor points (Fig. 2A, lower row).

**Figure 2:**
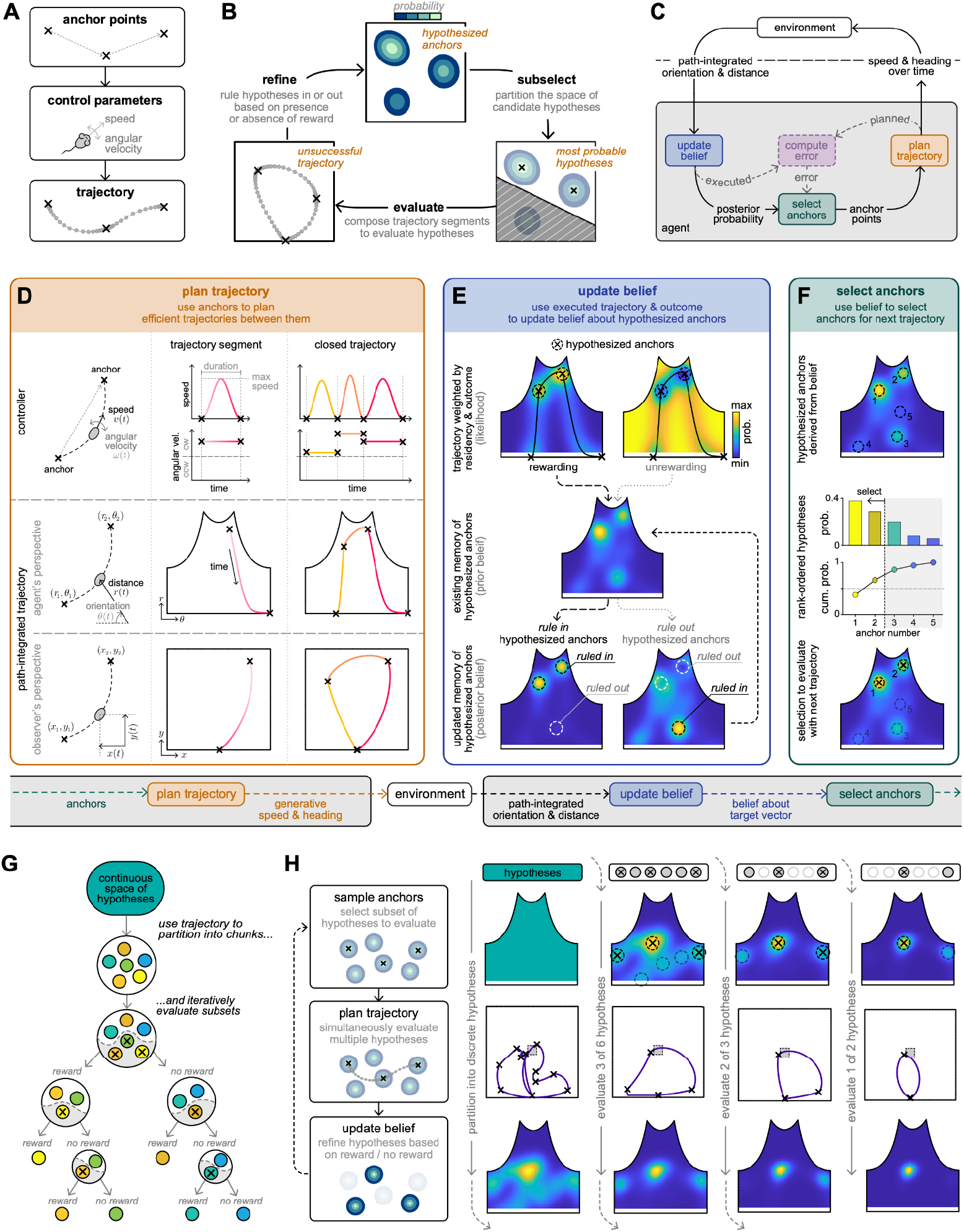
A minimal agent plans future trajectories based on the outcomes of past trajectories. **A)** We build an agent that generates trajectories via sets of ‘anchor points’. Vectors between pairs of anchor points are used to specify the parameters of a motor controller that generates smooth path segments between them. Together, a set of anchor points and path segments can be composed into trajectories. **B)** To guide the composition of successful trajectories, we assume that the agent maintains an internal belief about a successful anchor point. This belief can be used to specify a discrete set of hypothesized anchors (peaks in top panel); the agent then selects a subset of these hypotheses (lower right panel) to evaluate with its next trajectory (lower left panel). This trajectory, together with its outcome, is then used to refine the belief. **C)** We formalize the computations in panel B within three computational modules: a belief module tracks the agent’s belief about a successful anchor; a sampler module samples anchor points from the agent’s belief to guide future trajectories; the planner module plans and executes trajectories that intercept sampled anchor points. An additional error module can be used to adjust behavior based on differences between planned and executed trajectories. **D)** Planner module. The agent generates trajectory segments that traverse pairs of anchor points. To generate these segments, the agent controls its speed and angular velocity over time via a set of parameterized functions (upper row). These functions are fully specified by three parameters—the maximum speed, (constant) angular velocity, and total duration—that can in turn be derived, together with an initial heading direction, from the vector connecting any pair of anchor points (Methods). We assume that the agent tracks its path-integrated trajectory as a function of orientation and distance relative to the home port (middle row). This trajectory can also be visualized in cartesian coordinates from the viewpoint of an external observer (lower row). **E)** Belief module. The agent maintains an internal belief that specifies the probability that a candidate anchor will generate a successful trajectory. To update this belief, an executed trajectory is used to specify the likelihood of an observed outcome under different putative anchor placements (upper row). If a given trajectory was rewarded, probability mass will be concentrated along the trajectory (upper left); if the same trajectory was unrewarded, probability mass will be concentrated away from the trajectory (upper right). This likelihood is then combined with a prior belief (middle row) to construct an updated posterior belief (lower row); note that the two peaks near the top of the probability distribution are either strengthened (“ruled in”) or weakened (“ruled out”) depending on whether the trajectory was rewarded). **F)** Sampler module. The agent uses its internal belief to select anchor points to guide future trajectories. Peaks in the belief encode hypothesized anchors that the agent believes are likely to generate rewarding trajectories. The agent selects the most promising of these hypothesized anchors to evaluate on its next trajectory. To this end, the agent rank-orders the set of peaks by their probability, and selects the minimum number to cover 50% of peak distribution (a proxy for the full belief distribution; see Supplementary Methods). These selected peaks serve as anchor points for the next trajectory. **G)** When coupled together, these modules enable agents to rapidly narrow a space of hypotheses by first breaking a continuous space into discrete hypotheses, and then iteratively ruling them in or out. **H)** To localize hypotheses, the sampler selects hypotheses to evaluate (top row); the planner determines a trajectory that can evaluate all selected hypotheses with a single run (middle row); and the belief module refines the set of valid hypotheses based on the outcome of the executed trajectory (bottom row). Shown for a single example agent.

With this formulation, the problem of guiding behavior can be decomposed into the problem of appropriately selecting anchor points, and appropriately planning trajectory segments to intercept them. To jointly solve these problems, we construct an agent that tries to gather rewards while reducing behavioral expenditure, and thus tries to stabilize a successful single-anchor trajectory. To guide the selection of anchor points, we assume that the agent maintains an internal belief over a set of hypothesized anchor points that could yield successful trajectories; local peaks in this belief can be viewed as discrete hypotheses about the successful anchor. To rapidly localize within this hypothesis space, the agent selects a subset of these discrete hypotheses—in the form of anchor points—to evaluate with its next trajectory. It then executes this trajectory, observes the resulting outcome, and uses this information to refine its belief (Fig. 2B).

We formalize these computations within three distinct computational modules (Fig. 2C): (1) a belief module uses Bayesian inference to maintain and update a belief about the successful anchor point; (2) a sampler module selects among a set of hypothesized anchor points that are derived from the belief; and (3) a planner module plans a trajectory to evaluate the set of hypothesized anchors. Together, these modules comprise a behavioral policy that guides the agent’s future behavior based on the consequences of past behavior. Fig. 2D-F details how each of these modules is implemented, beginning with the trajectory planner.

To generate extended trajectories, we assume that the agent represents anchor points as vectors relative to a fixed reference, and that it controls its speed and angular velocity over time to sequentially intercept anchor points. We exploit the observed structure in mouse behavior by assuming that the agent speeds up and slows down between anchors while maintaining a constant angular velocity (Fig. 2D, upper row); the latter gives rise to linear changes in heading, as observed in mice (Fig. 1D). These generative functions are fully specified by the maximum speed, angular velocity, and total duration of the trajectory segment, which are in turn specified by the vector between two anchor points (Methods). More complex trajectories can then be generated by composing multiple segments in order to sequentially intercept a sequence of anchors. When integrated over time relative to a given initial heading, this results in smooth, curvilinear trajectories through space (Fig. 2D, middle and lower rows).

When executing a trajectory, we assume that the agent can track its path-integrated orientation and distance relative to the home port^4,23^. This information can be combined with a memory of hypothesized anchor points to rule in (if rewarded) or rule out (if not rewarded) candidate anchor points for guiding future trajectories (Fig. 2B). We formalize this within the context of Bayesian inference by iteratively updating an internal belief about the successful anchor point that generates a rewarding trajectory. On each trial, we use the executed trajectory and its outcome to construct a likelihood function that specifies how likely the observed outcome would be under different putative anchor placements (Fig. 2E, upper row). This formalizes the idea that if a trajectory resulted in a rewarding outcome, the successful anchor is likely to be along the executed trajectory; conversely, if the same trajectory resulted in an unrewarding outcome, the successful anchor is unlikely to be along the executed trajectory. In constructing this likelihood, we assign probability in proportion to residency; because the agent speeds up and slows down between anchor points, this leads to a relative increase (if rewarded) or decrease (if not rewarded) of probability mass at anchor points along the executed trajectory (Fig. 2E, upper row). The likelihood function is then combined with a prior belief about the successful anchor to construct an updated posterior belief (Fig. 2E, middle and lower rows).

Due to the spatiotemporal structure of navigational trajectories, the belief update described above will often lead to a multi-peaked posterior distribution, with multiple localized concentrations of probability mass. As described above, these local peaks can be viewed as discrete hypotheses about which anchor point will generate a successful trajectory. Given a set of hypotheses with differing probabilities, the fastest way to localize among them is to evaluate the smallest subset that covers 50% of the probability distribution, and to rule this subset in or out based on the presence or absence of reward. This approach maximizes expected information gain—such that rewarding and unrewarding outcomes are equally informative—and can be approximately achieved by selecting the fewest number of peaks in the belief distribution that cover 50% of the posterior probability spanned by the peaks (Fig. 2F, middle and lower rows; Methods). To evaluate this set of selected hypotheses, we assume that the agent guides trajectories through them. If a given trajectory is rewarded, the agent rules in the selected hypotheses, and rules out the remaining ones; conversely, if the trajectory is unrewarded, the agent rules out the selected hypotheses, and rules in the remaining ones (Fig. 2E, lower row). This is naturally implemented by the belief update, and allows the agent to learn equally well from rewarding and unrewarding outcomes.

When coupled together, these three modules interact to rapidly generate and modify compact trajectories that are consistent with reward (Fig. 2G-H). We assume that agents do not have any initial spatial biases when they enter the arena; this is captured by a uniform prior belief that encodes a continuum of equally valid hypotheses about successful anchors. Upon randomly sampling an initial set of anchors, agents partition this continuous space into discrete hypotheses, and iteratively evaluate subsets of these hypotheses (Fig. 2G). This is achieved by updating the internal belief based on the consequence of the previous trajectory, and using the updated belief to sample hypotheses that guide the subsequent trajectory (Fig. 2H).

Within tens of trials, agents reliably converge on single-anchor trajectories that intercept the target (Fig. 3A, left column). This rapid convergence is evidenced by an increase in reward probability (Fig. 3A, upper left) and is supported by the evolving structure of the internal belief. As agents refine their hypotheses about successful anchors, this belief becomes more localized over time, with fewer peaks and a lower overall entropy (Fig. 3A, middle left). Because anchors are sampled from this belief, the reduction in entropy is correlated with a reduction in the number of anchors along the trajectory, and thus with a reduction in the overall path length (Fig. 3A, lower left).

**Figure 3:**
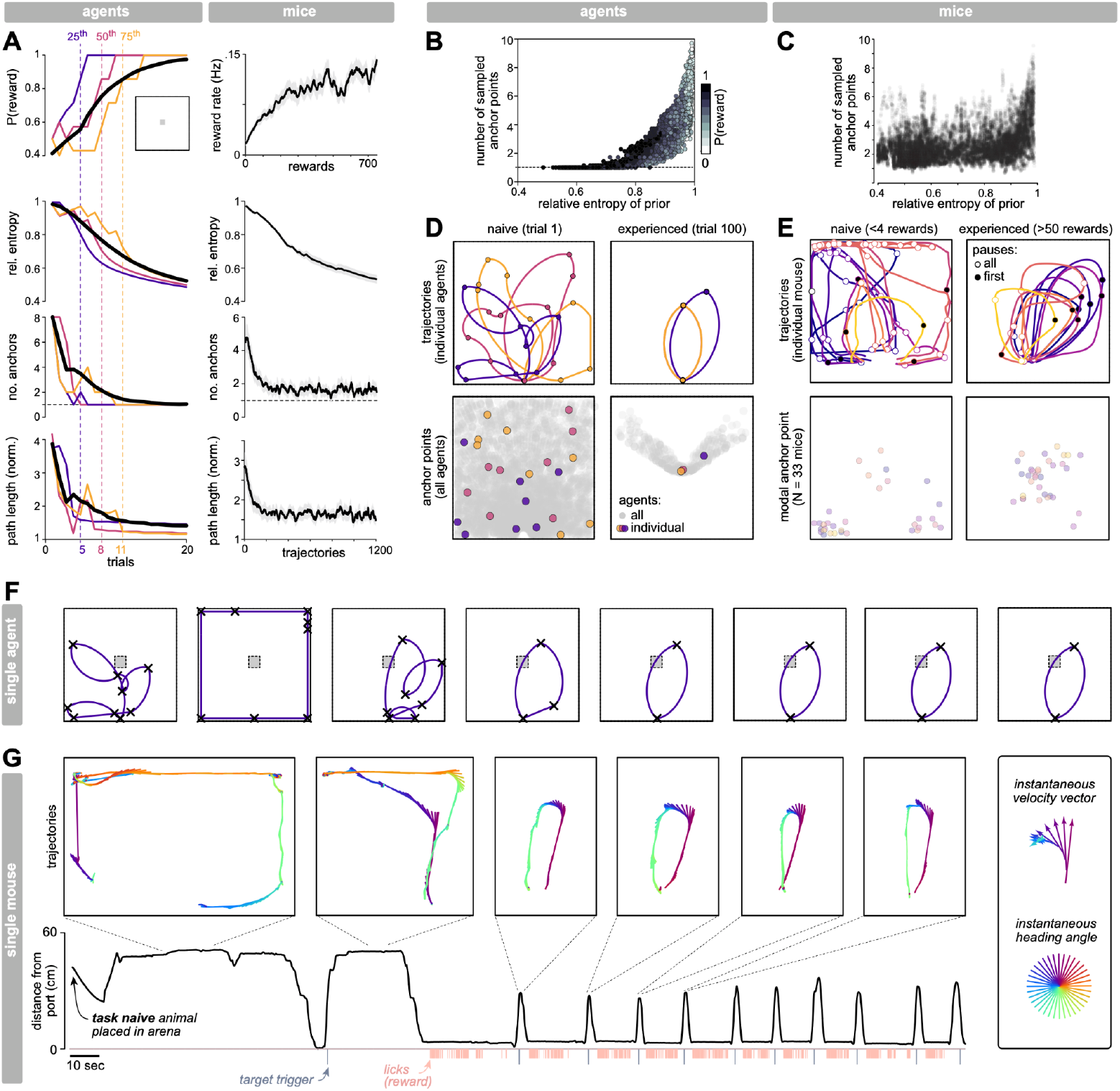
Agents and mice rapidly converge on compact trajectories that yield reward. **A)** When learning to intercept a hidden target placed in the middle of an arena (upper left inset), the behavior of agents (left column) rapidly evolves over time in a similar manner to that of mice (right column; top and bottom panels were reproduced from Fig. 1D). Agents learn to intercept the target within tens of trials; this is marked by a rapid increase in performance, decrease in the entropy of the posterior belief (measured relative to a flat distribution), and decrease in the number of anchor points and path length of trajectories (note that here and elsewhere, we exclude the two home anchors when reporting the number of anchor points). Black traces: average performance across 500 agents. Colored traces: three individual agents that reached reward probabilities of 6/7 within 5 trials (purple; 25th percentile), 8 trials (orange; 50th percentile), and 11 trials (yellow; 75th percentile). **B-C)** Learning is characterized by a structured relationship between the entropy of the prior belief before executing a trajectory—which is built from the history of past trajectories and their outcomes—and the number of anchor points that are sampled during the execution of a new trajectory. Shown for agents (B) and mice (C). **D)** Naive agents generate trajectories whose anchor points are spread across the entire arena (left column); after learning, anchor points are localized in regions that lead to consistent interceptions of the target (right column). **E)** Upper row: data from an example mouse, showing individual rewarded trajectories from the first session (upper left) and the 7th session (after ~50 rewarded trajectories; upper right). Filled dots indicate the end of the first inferred trajectory segment; open circles indicate subsequent segment boundaries. Anchor points are inferred from the spatial density of the first trajectory segment termini (see Methods for details). Lower row: for each of 33 mice, the modal anchor point was inferred from the histogram of first pause locations for naive trajectories (<4 rewards in first session, upper) and experienced trajectories (>5 sessions, bottom). **F)** Evolution of trajectories for the first agent shown in A, which reached criterion performance in 5 trials. **G)** Data plotted from the mouse that exhibited the fastest learning within our datasets (>30 subjects). Note that the mouse localized the target by the third trajectory, and continued to successfully intercept it thereafter. The lower trace shows the radial distance from the reward port, plotted in continuous time from the beginning of recording in the first task session. Red lines indicate the logic signal for detected licks at the reward port. Thin black lines indicate when the animal occupied the hidden target. Upper insets show extracted trajectories, defined by the moment the mouse left the reward port area (using a distance threshold) until it returned. “Feather plots” indicate the instantaneous heading vector angle (color) and speed (proportional to vector length).

The agent’s behavior captures the increase in reward rate and reduction in path length observed in mice over the course of learning (Fig 3A, upper and lower right; also shown in Fig 1D). However, this formulation additionally predicts that the number of anchor points on a given trial should correlate with the belief built from the trajectories executed on previous trials (Fig. 3B). This prompted us to use sequences of mouse trajectories to infer an evolving posterior belief, and assess the entropy of that belief over time and in relation to the boundaries between trajectory segments (i.e., putative anchor points). As predicted, both the entropy of the belief and the number of putative anchor points decrease over time during learning (Fig. 3A, middle right) and are correlated with one another on a trial-to-trial basis (Fig. 3C).

This behavior of agents reveals a key property of trajectory learning: agents stabilize trajectories that consistently yield reward, but they do not learn the target location *per se*. This can be observed in the distribution of anchors as they stabilize over learning and extended performance (Fig. 3D). Before learning, agent trajectories are composed of multiple anchor points that are distributed across the arena (Fig. 3D, left column); after learning, their anchors are localized in regions that enable reliable interception of the target (Fig. 3D, right column). This prompted us to analyze the spatial distribution of putative anchor points in mice. When mice are naive to the task or experience a sudden change in target location, their initial trajectories are composed of multiple segments whose putative anchors are broadly distributed around the arena (Fig. 3E, left column; see Methods for details). With experience, these putative anchors get localized to the vicinity of the hidden target (Fig. 3E; right column). As with agents, mice stabilize trajectories that consistently yield reward, and need not learn the target location *per se*.

Notably, the fastest agents learn in one shot, and are able to reliably intercept targets after receiving a single reward. This requires a fortuitous sequence of trajectories, and thus happens infrequently (Fig. 3F); nevertheless, this predicts that the occasional mouse might also approach such limits. Indeed, when we examined learning trajectories across individual mice, we observed an example of this exceptionally rapid learning (Fig. 3G): the fastest mouse in this cohort (>30 animals) localizes the target within three trajectories (or 1-2 reinforcements), and continues to reliably intercept the target thereafter.

When taken together, this algorithmic formulation captures the evolution of many different aspects of behavioral structure over the course of learning, and simultaneously captures the upper limits of learning efficiency observed in animals. There are several key factors that enable this rapid convergence; SI Fig. 1B illustrates these factors in comparison to a more canonical tabular representation (SI Fig. 1A) that is imbued with the same core capabilities of our algorithm. First, behavioral trajectories are fully specified by a compact set of control parameters that can be derived from vectors between anchor points. For example, a three-segment trajectory is specified by seven control parameters (SI Fig. 1B-i), which is far fewer than is required if specifying the coordinates of individual points along the trajectory or elemental steps on a predefined grid (SI Fig. 1A-i). This compact set of control parameters substantially reduces the cost of planning relative to a larger set of parameters. Moreover, because these control parameters shift and scale a set of generative functions, the resulting trajectory is structured in both space and time. This structure is exploited by the likelihood function within the Bayesian update, and is important for partitioning a continuous space of hypotheses into discrete chunks (SI Fig. 1B-ii). This partitioning is not achieved when specifying trajectories as steps on a grid, where all steps have equal weight (SI Fig. 1A-ii).

The advantages of this partitioning are most notably seen when considering a flat prior belief that encodes a continuum of hypotheses about successful anchors; after executing a single structured trajectory, the updated posterior belief has multiple peaks and valleys concentrated around places where the agent slowed down. These peaks provide a natural chunking of the belief space, and enable the agent to easily prioritize and sample hypotheses within this space. The agent uses the outcomes of its trajectories to evaluate these hypotheses, and in doing so takes advantage of both rewarding and unrewarding outcomes. As a result, the agent can learn just as much from the absence of reward as it can from the presence of one (SI Fig. 1B-iii), which speeds learning relative to using reward alone. To efficiently narrow down within this hypothesis space, the agent selects hypotheses based on their prior probability, which provides an unambiguous method for prioritizing some hypotheses over others (SI Fig. 1B-iv). Without a chunking of the hypothesis space, there are many possible hypotheses that are equally valued, and many possible ways to partition among them (SI Fig. 1A-iv). Finally, because the hypotheses themselves fully specify the control parameters of the trajectory planner, planning is straightforward (SI Fig. 1B-v). This is not typically the case in tabular representations, where one has to specify many additional parameters to traverse a given set of hypothesized grid locations (SI Fig. 1A-v). Thus, all three modules work together to exploit the structure of behavior and thereby enable rapid learning.

### Internal beliefs can be used to assess errors and adapt to environmental changes

The previous results highlight how an internal belief, in combination with a generative model of behavior, can be used to guide actions that yield successful outcomes and strengthen existing beliefs. However, these modules can also be used to interpret unsuccessful outcomes that are inconsistent with existing beliefs. We next explored two different scenarios—obstacles and moving targets—that animals encounter in navigational settings, and that introduce deviations between internal beliefs and external outcomes. In both cases, we find that mice rapidly adapt trajectories in this hidden target paradigm, and we show how internal beliefs can be used to assess errors and modify behavior in a manner that captures observed patterns of behavioral flexibility.

#### Execution errors enable rapid reconfiguration of trajectories around obstacles

When the configuration of the environment changes—for example, when an obstacle is introduced—animals might need to divert their previously-planned trajectories in order to navigate to existing goals. This diversion provides localizing feedback about how planned trajectories should be modified to more efficiently avoid the obstacle in the future. To illustrate this, consider an agent that diverts its trajectory around a newly encountered obstacle before resuming a planned path. This diversion manifests as an error between the planned and executed paths, and is localized to the region occupied by the obstacle (Fig 4A). Tracking this error relies on the same core computations that we introduced above; indeed, if we compare the path-integrated planned and executed trajectories, their difference provides a local, directional error signal that differentiates successful from unsuccessful segments of the trajectory. Information about those segments that were successful in the past can then guide trajectory segments that will be successful in the future.

**Figure 4:**
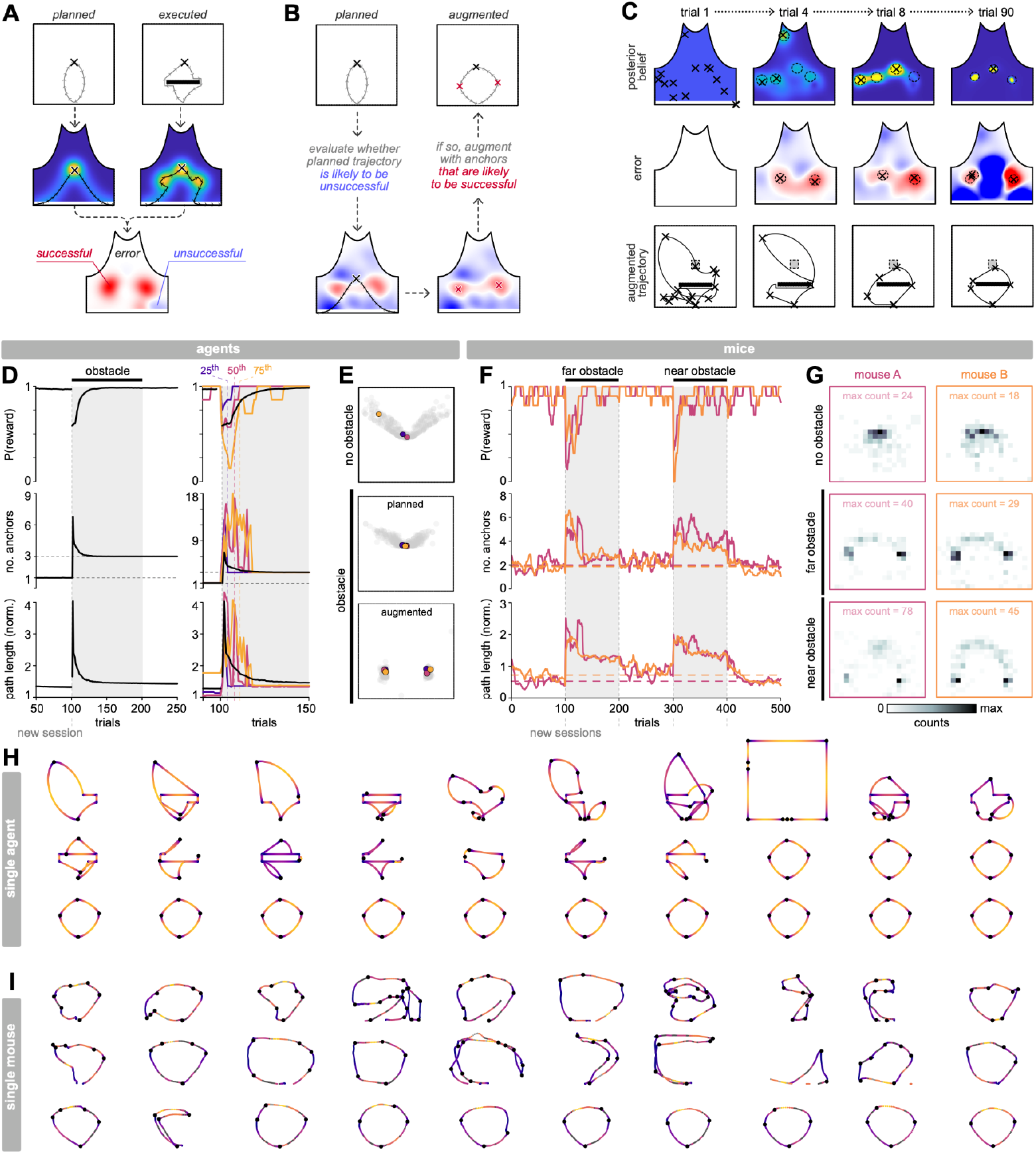
Path-integrated execution errors can guide obstacle avoidance. **A)** When an obstacle is introduced into the arena, the agent must divert its trajectory around it. This leads to differences between the planned (left column) and executed (right column) trajectories. The difference between the path-integrated trajectories can be used as an error function that differentiates portions of the planned trajectory that could not be executed (blue; ‘unsuccessful’) from portions of the executed trajectory that successfully diverted the agent around the obstacle (red; ‘successful’). **B)** When planning a new trajectory (upper left), the agent can use the error function to predict whether its planned trajectory will pass through regions of the arena where previously-planned trajectories could not be executed (blue regions of heatmap; lower left). In such cases where the planned trajectory generates high predicted error, the agent can use this same error function to select additional anchor points (red x’s, lower right) to ‘augment’ its trajectory; this enables the agent to more efficiently avoid obstacles (upper right). **C)** The agent’s target belief and target error co-evolve in a self-consistent manner during learning. Both the target belief (first row) and the target error (second row) are used to select anchor points to guide trajectories (third row). Over time, the agent learns a self-consistent set of 3-4 anchor points that allow it to intercept the target while avoiding the obstacle (last column). **D)** Behavioral responses to obstacles, averaged across agents (left column; 500 agents) and shown for three individual agents (right column) that reached reward probabilities of 6/7 within 4 trials (purple; 25th percentile), 8 trials (orange; 50th percentile), and 11 trials (yellow; 75th percentile) after the obstacle was introduced. Agents first learn to intercept a target in the absence of an obstacle. We assume that the obstacle is introduced in a new session, for which agents begin with a flat belief; just after the obstacle is introduced, agents show a transient drop in reward probability (upper row), and a transient increase in the number of anchors (middle row) and the overall path length (lower row) of their trajectories. Within tens of trials, agents refine their trajectories and recover high reward probabilities. **E)** Relative to conditions with no obstacles (upper panel), agents tend to stabilize trajectories that use one anchor point to intercept the target (derived from the posterior belief; middle panel), and two additional anchor points that avoid the obstacle (derived from path-integrated errors in previous trajectories; lower panel). **F-G)** Same as D-E, but shown for two individual mice (pink and orange curves) that each experienced two different obstacles introduced at the start of new sessions (Methods; only first 100 trajectories around session starts are plotted). Mice show similar changes in reward probability, number of putative anchors, and path length after an obstacle is introduced. Histograms in G were computed over 50 trials for each mouse. **H-I)** Evolution of trajectories for an individual agent (H) and individual mouse (I), shown for 30 successive trials following the introduction of an obstacle. Black dots indicate anchor points. Color indicates relative speed along the trajectory, with purple and yellow indicating lower and higher speeds, respectively.

To exploit this knowledge, the agent can integrate this error signal over time and use it to evaluate the predicted error along a planned trajectory. If the planned trajectory passes through regions along which trajectories had previously been diverted—i.e, if the planned trajectory is predicted to generate large execution errors—the agent can use this same error signal to sample new anchors to augment its planned trajectory (Fig. 4B). These anchors are localized to regions where the agent has made errors in the past, and thus enable the agent to augment its trajectory in order to minimize errors in the future. Fig. 4C shows how the anchor belief, error, and trajectories co-evolve in a single agent that must simultaneously localize a new target while avoiding an obstacle. To minimize predicted errors in its trajectories, the agent stabilizes anchor points at the far edges of the obstacle, in addition to an anchor point that enables the agent to intercept the target.

This ability to augment trajectories with new anchor points captures a variety of different behavioral patterns observed in mice (Fig. 4D-I)^7^. When agents and mice first encounter an obstacle after having previously learned to intercept a hidden target, both show a transient drop in reward probability and a transient increase in the path length and number of anchor points along their trajectories (Fig. 4D,F). Within tens of trials, they rapidly recover their earlier performance while stabilizing compact trajectories (Fig. 4H,I) that include two anchor points at the edges of the obstacle (Fig. 4E,G). Mice reliably exhibit these patterns for two different obstacle placements introduced separately in time (‘near’ and ‘far’ obstacles in Fig. 4F,G; see Methods for details). Once the obstacle is removed, both agents and mice initially continue to intercept previously-learned anchor points, but mice more quickly relax to trajectories with shorter path lengths and single pauses. This hints at the possibility that the obstacle removal could provide additional cues that trigger a resetting of the error signal and a corresponding trimming of anchors along the trajectory.

#### Surprising outcomes enable memory caching of behavioral policies to support rapid contextual inference

The previous results highlight how changes in the configuration of the environment provide localizing error signals that can be used to modify behavioral trajectories. When changes are made to the location of hidden targets, however, there are no localizing cues available to the agent to signal the change. Instead, the agent can use its internal belief to predict the probability of expected outcomes, and detect violations of these predictions.

When the agent has learned to reliably intercept a target, its belief is localized in a manner that generates rewarding trajectories. As a result, the agent predicts reward with high probability, and is unsurprised when the outcome of its trajectory is consistent with that prediction (Fig 5A, left column). However, if the target location abruptly changes, the agent still predicts reward with high probability, but the same trajectory no longer yields reward. This results in a mismatch between the agent’s internal prediction and the observed outcome, and yields high surprise (Fig 5A, right column). This mismatch is accompanied by a prolonged drop in reward probability as the agent slowly updates its old belief (Fig. 5B). Rather than suffering through this prolonged period of slow learning, the agent can instead exploit the surprising observation and use it to trigger a resetting of its belief (Fig 5C). When encountering new target locations (Fig. 5D, upper row), this resetting enables agents to intercept different targets equally fast, regardless of their spatial separation (Fig. 5D, lower left). Mice behave similarly; when repeatedly exposed to new target locations, mice can localize targets equally fast regardless of their separation, and in many cases achieve the same high learning efficiency as agents (Fig. 5D, lower right; shown for the fastest sessions).

**Figure 5:**
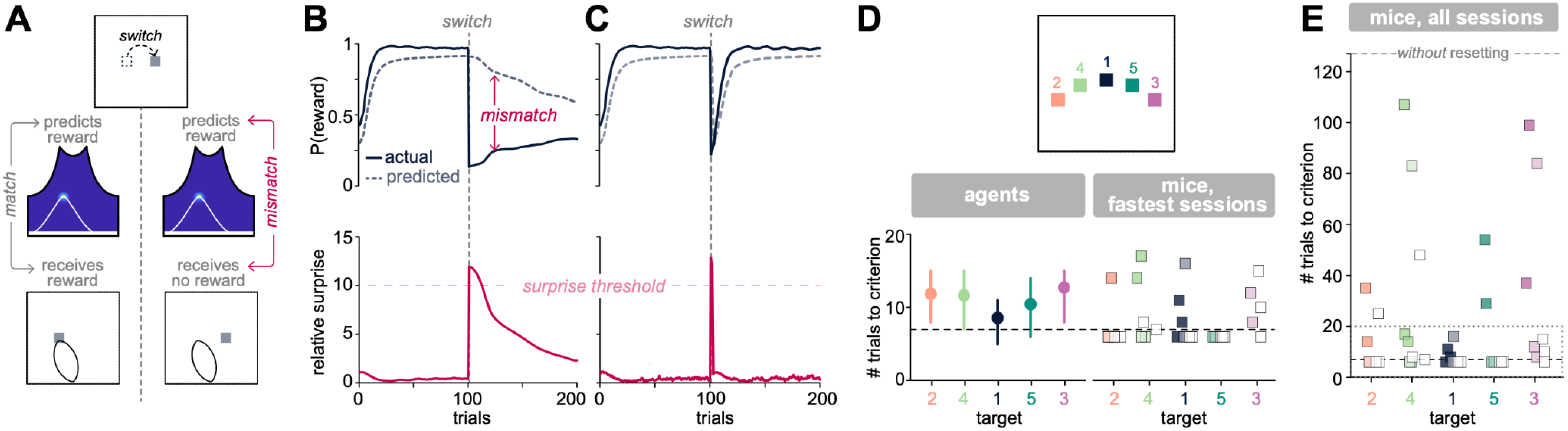
Violations of internal beliefs can signal context changes. **A)** Left column: after learning, the agent maintains a localized belief that generates rewarding trajectories. As a result, the agent’s prediction of reward when planning trajectories is matched to the outcome that it receives upon executing those trajectories. Right column: when the target switches locations, the agent continues to predict that its planned trajectories will yield reward; however, when executed, those trajectories no longer intercept the new target. This leads to a mismatch between predicted and actual reward. **B)** Upper: as schematized in panel A, a target switch leads to a prolonged mismatch between predicted and actual reward rates. Lower: as a result of this mismatch, the agent is surprised about the outcome of its trajectories; this high surprise arises because incoming observations are inconsistent with the agent’s belief. When this surprise is sufficiently high (e.g., when it exceeds a fixed threshold), it can signal a change in the environment and can be used to reset the agent’s prior belief. **C)** When agents are allowed to reset their prior belief following a sufficiently surprising outcome, they quickly localize a new target after a switch. The red horizontal dashed line marks the surprise threshold used to trigger a belief reset. Results in B and C were averaged across 500 agents. **D)** Upper: Schematic of five targets that were presented sequentially in time. Lower left: Agents localize targets equally quickly, regardless of their spatial separation. Circular markers denote the average number of trials required to reach a criterion reward probability ≥ 6/7, and solid vertical lines span the 25th to 75th percentile across 500 agents. Lower right: On the fastest sessions, mice localize targets as quickly as agents. Square markers denote learning speeds computed from individual sessions for each of three mice; the first session was excluded in cases where mice did not adapt to the target switch (Methods). The horizontal dashed line shows the 25th percentile across agents that were allowed to reset their beliefs. **E)** Across all sessions, the distribution of learning speeds in mice falls between agents that do and do not reset their prior beliefs. The lower and upper horizontal dashed lines show the 25th percentile across agents that were (lower) and were not (upper) allowed to reset their beliefs; the dashed box shows the sessions that were highlighted in D.

Not all mice show this rapid localization on all sessions; across individual sessions, the distribution of learning efficiency falls between those agents that do and do not reset their prior beliefs following sufficiently surprising outcomes (Fig. 5E). This variability suggests that mice might hold onto localized posterior beliefs after the environment has changed, while also exhibiting the rapid learning that arises from resetting prior beliefs. Indeed, after having refined an accurate posterior belief that reliably generates good outcomes, it can be advantageous to retain that belief in memory while simultaneously adapting to changes in the environment. Consistent with this observation, Figure 6A plots the endpoint of the first outward trajectory segment for two mice across multiple target switches. After a target switch, mice continue to sample anchors that are consistent with previous targets, and they also—in parallel during interleaved trajectories—rapidly acquire new anchors that are accurately directed towards the current target (Fig. 6A). Of those trajectories consistent with previous targets, many are directed toward the most recently rewarded target, but some are also directed toward targets encountered further in the past. This suggests that mice retain memories of multiple behavioral policies that had been successful in past environmental settings while simultaneously learning new behavioral policies that are appropriate for the current setting.

**Figure 6:**
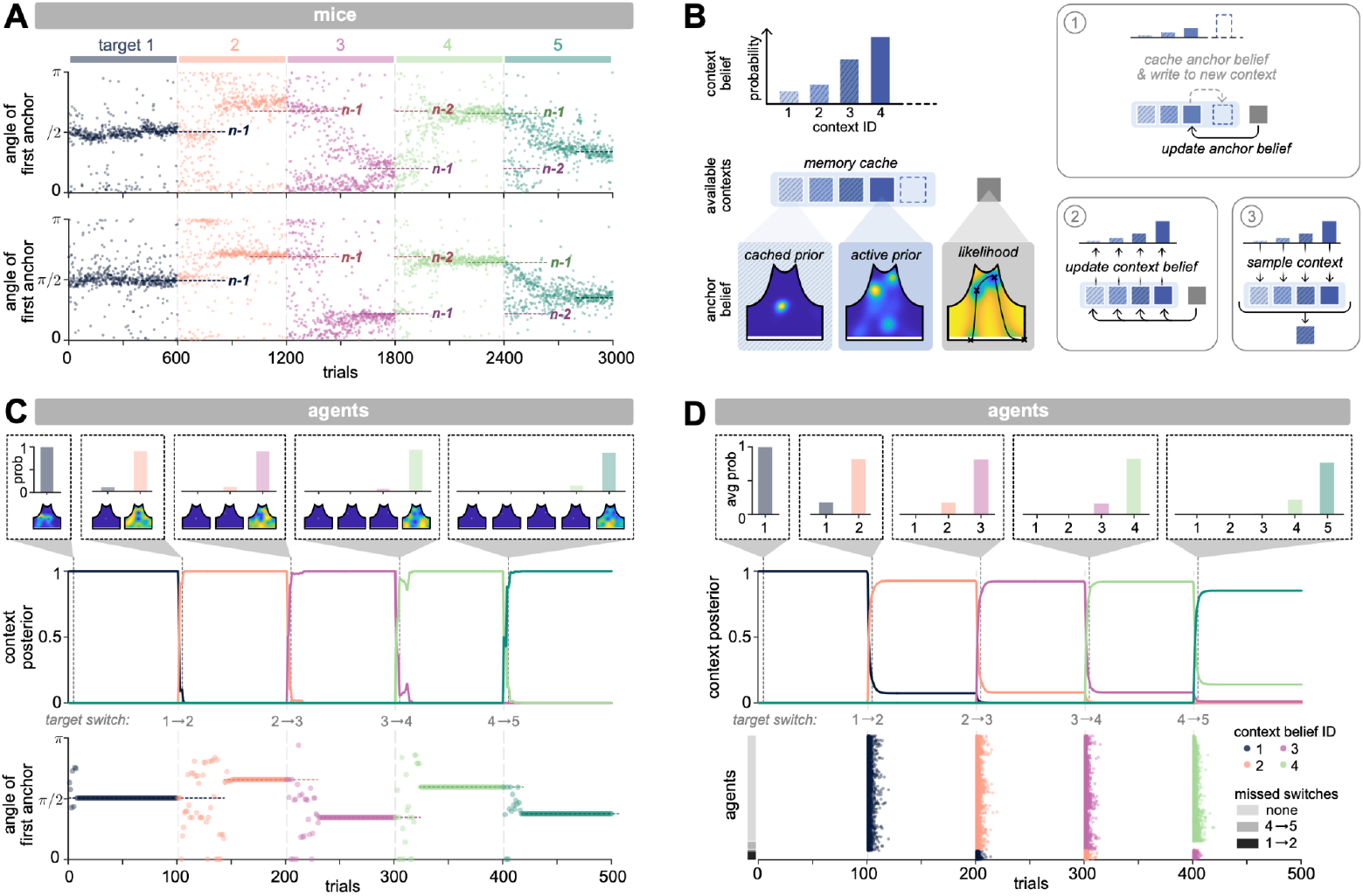
A flexible memory cache supports contextual inference. **A)** Angle to the endpoint of the first trajectory segment (i.e., to the first anchor point), shown for two individual mice across multiple target switches. After a target switch, mice simultaneously acquire new anchors while intercepting anchors that were previously rewarding. In a given block, mice often sample anchors that were rewarding in the previous block (‘n-1’), or in two blocks prior (‘n-2’). **B)** When the environment changes, it can be advantageous to retain the current belief in memory to be used again in the future. We allow the agent to maintain these past beliefs in a memory cache; these beliefs serve as internal models of different contexts that the environment can occupy. Box 1: To maintain and use this cache, the agent uses its executed trajectory and outcome to update a belief about the successful anchor within a context; if the outcome is sufficiently surprising under the current belief, the agent caches the current belief and begins updating a new belief within its cache. Box 2: The agent uses its trajectory and outcome to update a belief about current and past contexts. Box 3: This context belief is used to sample candidate contexts (and their corresponding anchor beliefs) to guide future trajectories. **C)** When an individual agent is allowed to cache and recall previously-stored beliefs, it begins with a belief that there is only one environmental context (black curve in upper row), and updates its belief about the successful anchor within that context. After the first target switch, the agent is sufficiently surprised by the outcome of its trajectory under its current anchor belief; it caches that belief and begins writing to a new anchor belief under a new context (orange curve in upper row). Within a few trials, the agent believes that this new context best explains the outcomes of its trajectories, as indicated by the orange curve approaching a probability of 1. During this time, the previous anchor belief remains cached, but can be sampled on individual trials. This is evidenced by the agent making directed runs to previously-learned targets (lower row; note the orange points near the orientation of the previous target). This behavior diminishes over time as the agent learns to localize new targets and appropriately update its context belief. This behavior continues for each new target switch as the agent iteratively grows its context belief. **D)** On average, agents exhibit similar behavior to the single agent shown in **C**. Upper row and insets: context posterior averaged across 500 agents. Lower row: times when individual agents sample from previously-cached posteriors (labeled by their context ID). Note that some agents do not detect all target switches.

This behavior naturally arises if agents are allowed to store their posterior belief in memory when the environment changes, and reuse this belief in future settings. Indeed, the abrupt increase in surprise that follows a target switch can be used not only to reset the agent’s belief, but also to expand its scope, for example by allowing for the possibility of different environmental contexts that necessitate different behaviors. A surprising outcome can thus be interpreted as a change in context, and can be used to prompt the agent to commit its current belief to memory before resetting its prior. This ‘cached’ belief stores a behavioral policy that was successful in a particular context, and that could be useful again in the future.

Iteratively caching beliefs has the advantage of enabling the agent to grow its internal model of the world based on experience, rather than assuming a fixed internal model. However, the agent must then decide how to weigh its current belief against those maintained in its memory cache (Fig 6B). This is naturally accommodated within the same Bayesian formalism by building and updating a belief *about the current context*, which is then used to query a belief *about the successful anchor point* within a context. Upon executing a trajectory, the agent evaluates the probability of the resulting outcome under both its current and previously cached anchor beliefs, which is in turn used to update the context belief.

When agents are allowed to cache and recall previously-held beliefs, they make directed runs to previously-used anchor points (Fig 6C-D), much like mice. This diminishes over time as agents update their belief about the current context, and about the successful anchor within the current context. In settings where targets are never reused, this slows learning; however, if targets are reused, the memory cache allows agents to quickly recall previous beliefs without having to relearn from scratch.

## Discussion

We developed an algorithmic formulation of animal learning that modifies parameters of memory-guided navigational trajectories in order to achieve desired consequences, such as collecting rewards that are sparsely delivered and delayed in time. Our algorithm uses previous trajectories and their outcomes to update a posterior belief that informs the generation and composition of future trajectories. This belief evolves over time based on the consistency of past trajectories and outcomes, and results in a natural progression from highly structured exploration to targeted exploitation. This, in turn, enables exceptionally rapid learning that matches the upper limits of learning efficiency seen in mice, in contrast to previous approaches that are orders of magnitude less efficient than animal learning.

This rapid flexibility is supported by a compact generative model that uses a set of vectors to specify motor control parameters. Thus, vector-based computations are particularly germane to the problem of generating memory-guided navigational trajectories. Moreover, several lines of evidence suggest that activity in the hippocampus and associated cortical areas represents both past trajectories^5,24–27^ and vectors to task-relevant extrapersonal targets^28–32^, thereby hinting at biologically plausible implementations of this algorithm. What about allocentric, map-like strategies that have strong empirical support in brain activity and are thought to be critical for spatial learning? First, we note that while our agent represents anchor points as vectors relative to a fixed reference, this does not limit the agent to an ‘egocentric’ strategy; *i.e*., one that always guides behavior in a given direction relative to a new starting point^33^ (Fig. 7A). Rather, our agent can readily employ ‘allocentric’ strategies that intercept stored anchor points from new starting locations within the arena (Fig. 7B). This is naturally implemented by our trajectory planner, which uses vector subtraction—for which there are known neural mechanisms^34–37^—to compute relative distances between anchor points and specify appropriate control parameters. This in turn enables the agent to immediately intercept previously-learned anchor points from new starting locations within the arena (Fig. 7C). We have observed some evidence of such behavior in mice. For example, after long pauses at different locations in the arena, mice make directed runs that efficiently intercept the target on their way back to the home port (Fig. 7D).

**Figure 7:**
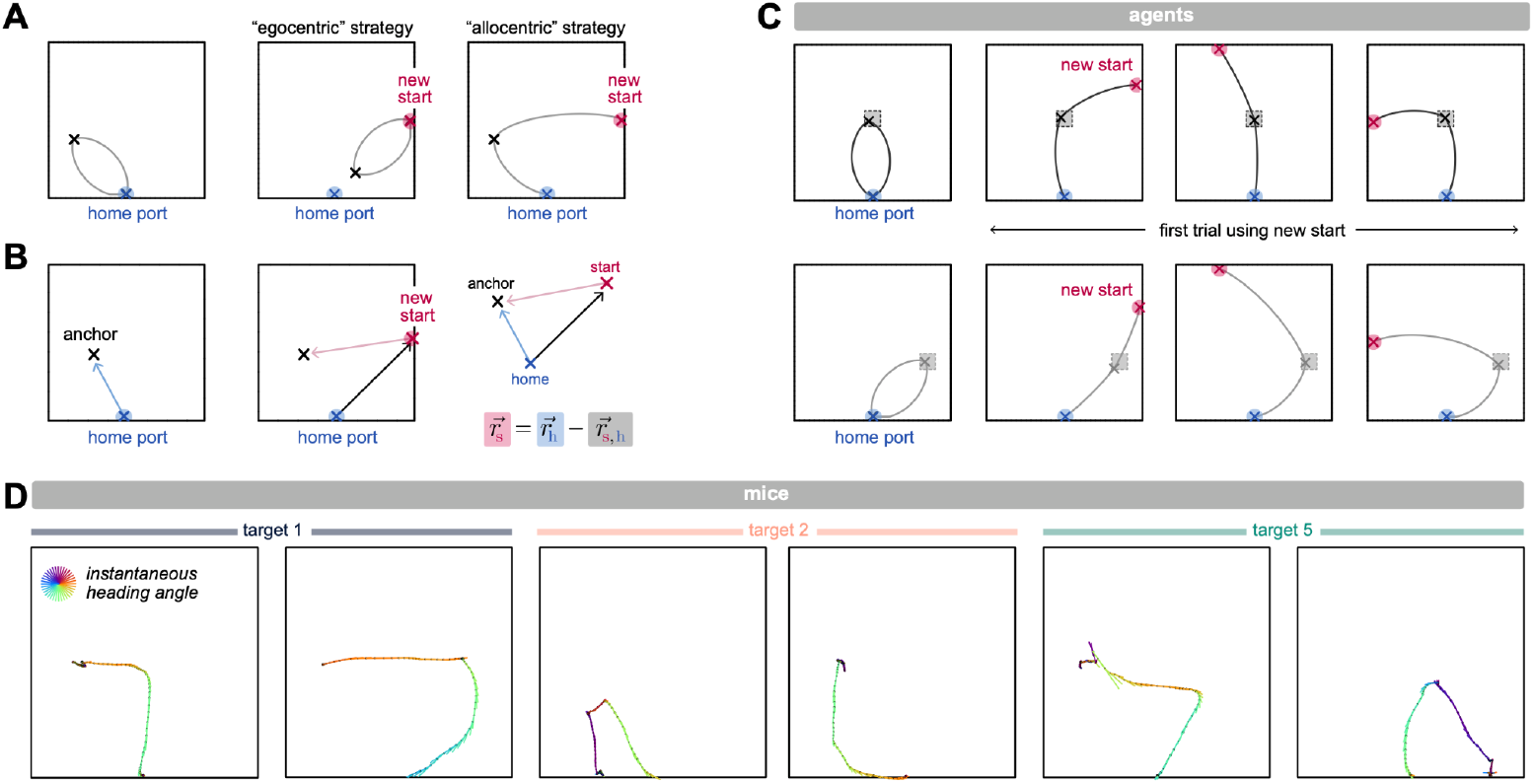
Trajectory learning enables flexible navigation to allocentric goals. **A)** A navigational trajectory can support an ‘egocentric’ strategy, in which an animal travels in the same direction relative to its starting location (left and middle panels), or an ‘allocentric’ strategy, in which it travels to the same location in allocentric coordinates (left and right panels). **B)** Our agent stores anchors as vectors defined relative to a fixed reference (here, the home port), and need only specify the vector distance between anchor points to control its speed and angular velocity over time. Thus, when the agent enters the environment from a new location, it can perform vector subtraction to compute the anchor vector relative to its starting location, and can use this to specify the necessary control parameters for navigating from the starting location to the remembered anchor. **C)** When agents enter the arena from a new entrance, they can immediately specify the correct control parameters to intercept anchor points that are referenced relative to the home port (shown for two different target locations). **D)** Instances where mice, after making long pauses at a large radial distance from the reward port, initiated rapid, directed runs that intercept the target on their way back to the home port. Trajectories are represented as in Fig. 3G.

Map-like strategies have an additional feature that provides an important and potentially critical complement to the trajectory-based algorithm described here. Specifically, maps provide a useful construct for formalizing knowledge about the extrapersonal, spatiotemporal structure of the environment that is critical to many kinds of spatial memory and learning. These constructs can be integrated with the algorithms described here, for example by layering ‘scaffold’-like representations onto navigational trajectories. As such, the ability to generate and rapidly reconfigure navigational trajectories can subserve a wide range of computations.

### Planning in the space of trajectory control parameters

To gather information from the environment, our agent plans and executes structured behavioral trajectories through space. Rather than solving an optimal path planning problem to generate these trajectories^38^, we looked to the structure of animal behavior to constrain an appropriate generative model. This prompted us to formulate a behavioral policy in terms of generative quantities that the brain controls^5,39,40^, rather than in terms of primitive actions or elemental transitions between a set of abstract states. The structure of behavior also suggested a hierarchical decomposition of trajectories into discrete segments delineated by pauses. This parametrization allowed us to reformulate the learning problem as one of selecting a compact set of putative anchor points that in turn specify the maximum speed, duration, and angular velocity along individual trajectory segments. The construction of trajectory segments as subpolicies is conceptually similar to the notion of ‘options’ within reinforcement learning^41^, but circumvents the problem of discovering state-action sequences that navigate to bottleneck or landmark states^42,43^. By tethering parameterized subpolicies to a compact set of anchor points, we substantially reduce the dimensionality of the planning problem. When compared to models that partition space into a grid of discrete states, and that guide behavior via sequences of elemental state transitions^10,14,15,44^, this formulation has far fewer parameters and learns much more quickly, capturing both the speed and evolving structure of animal behavior over the course of learning.

### Bayesian inference for refining hypotheses about anchors

To guide the composition of trajectories, the agent maintains a belief about the anchor point that will generate a successful trajectory. This enables the agent to appropriately assign credit to entire trajectories based on their consistency with a given outcome. To update this belief, the agent uses path-integration mechanisms to retain a memory of its most recent trajectory, and uses this memory to construct a likelihood function conditioned on whether or not that trajectory led to a reward. In constructing this likelihood, we assign probability based on time spent at different points along the trajectory; this has the computational advantage of partitioning a continuous belief space into discrete chunks—a property that is exploited by the sampler—and has the mechanistic advantage of being plausibly implementable by simple, Hebbian-like plasticity rules^40,45,46^. In the context of classic reinforcement learning algorithms, this memory is consistent with maintaining an eligibility trace^44,47^ that decays more slowly than the duration of typical trajectories. We further assume that when incoming observations are inconsistent with existing beliefs, the agent can cache its posterior belief to be reused in the future, as previously suggested for storing and reusing motor policies^48^. To query this anchor belief, we assume that the agent maintains a higher-order belief about environmental context, and can adaptively grow or shrink the set of plausible contexts over time. This has the advantage of enabling the agent to begin with a simple model of the task and augment as needed^49^, rather than assuming a fixed model *a priori*. More generally, this belief can be formulated to include explicit knowledge about other properties of the environment, such as target dynamics or other extrapersonal objects.

### Active sampling for narrowing the set of candidate hypotheses

The process of selecting anchors can be viewed as a form of active sampling, whereby the agent actively selects the data that it wishes to query in order to iteratively improve its model of the environment. We took an information theoretic approach and formulated this problem as a maximization of expected information gain^50–52^. Thus, the agent makes decisions to gather information—rather than to collect reward *per se*—and in doing so, rapidly achieves high reward rates. In the context of navigation, selecting appropriate data necessarily involves moving through space, which can require additional motor planning. Previous navigational algorithms, such as infotaxis^53^, address this by sequentially gathering new data through locally available actions. Here, we allowed the agent to gather data by planning a sequence of nonlocal actions, but we simplified the problem by separating the selection and motor planning processes. Rather than specifying the optimal trajectory that maximizes expected information gain across the entire belief space, we solved a heuristic approximation by selecting among local peaks in the belief space. This enabled us to transform a continuous selection problem into a simpler discrete one; the downstream planner then solves the separate problem of guiding trajectories through the set of selected points. In new environments, this leads to an initial exploration of space that is highly structured, and a gradual shift toward more exploitative behavior. This differs from typical reinforcement learning approaches that treat exploration and exploitation as distinct behavioral modes that are randomly interleaved over time^44,54,55^. Here, the degree of exploration emerges naturally through the internal belief, which leads to more exploratory behavior when the agent is uncertain and the belief is diffuse, and more exploitative behavior when the agent is certain and the belief is localized. Existing models of animal learning^5,39,56,57^ have addressed the exploration problem^54^ by injecting noise about previous solutions, which can be useful for detecting local target shifts without needing to revert to a uniform belief and random sampling of anchors. In the future, these two different modes of injecting variability could be combined to leverage the relative advantages of local and global exploration.

Although we assume that the selection of anchor points is driven primarily by maximizing expected information about a single successful anchor, there are in principle many different objectives that can be used to guide this selection. We considered one such objective when navigating around obstacles; here, the addition—rather than subtraction—of anchor points can be useful for diverting trajectories around obstructions. Rather than requiring the agent to build and store a map of obstacles and their spatial relations, we assume that the agent can use local error feedback about the success or failure of previously planned trajectories in order to adjust future plans. Path integration^23,58^ provides a natural mechanism for tracking this error, and the addition of new anchor points provides a natural mechanism for reducing this error. When multiple different objectives are used to drive the selection of anchor points, stable learning must converge on a self-consistent solution. Indeed, we found that agents converged on a set of stable anchor points that enabled them to simultaneously intercept the target while avoiding the obstacle. These stable anchor points can be interpreted as ‘subgoals’ that better enable the execution of a primary goal^7,14^.

### Generalization to more complex settings

The computational elements invoked here may be useful in more complex settings that require more advanced planning. Planning in large and partially unobservable state spaces is computationally costly and suffers from the curse of dimensionality. To combat this, different methods have been developed to perform tree searches over a reduced set of future states^59^ that can be prioritized based on their potential impact on planning^15^, or reuse planned policies to limit the need for replanning^60^. Here, we provide a computationally efficient approach for using a stored memory to iteratively trim a set of anchor points over which motor plans are constructed, and for storing previously-planned motor policies to be reused in the future. This is especially useful in settings where the environment is relatively stable over time. In sensory-driven settings, or in settings where motor plans have to be revised on faster timescales, these approaches could be combined with existing methods for identifying points of interest—for example, by computing the connectedness of space^15,61^—to guide motor plans based on a combination of sensory- and memory-driven inputs.

### Common building blocks of navigational learning across species

Navigation is critical across the animal kingdom, and studies in other species provide mechanistic support for several aspects of the computations that we invoke here. Recent work in insects has made substantial progress in understanding the neural mechanisms underlying flexible navigation^35,62–64^, and recent work in zebrafish suggests that many of these mechanisms might be shared with vertebrates^65^. For example, navigational learning studies in flies have identified candidate circuit architectures for building and updating a one-dimensional internal goal memory using an internal representation of head direction^35,40,66–68^, which is itself updated through the integration of self-motion signals^69,70^. This allows the goal memory to be updated even in the absence of any localizing sensory cues. This update is thought to be mediated by plasticity that modifies synapses based on the amount of time spent at different headings^35,40,66,67^, and recent physiology has shown how the resulting memory is used to guide steering commands toward a single goal heading derived from it^71,72^. Our algorithm relies on many of the same computational elements. For example, we assume that our agent maintains and updates a two-dimensional goal memory—in the form of a posterior belief—using an internal representation of orientation and distance that is itself updated through the integration of self-motions signals. We further assume—via the construction of our likelihood function—that this update depends on the time spent at different orientations and distances, similar to the residency-dependent plasticity thought to update the fly’s goal memory. Finally, we assume that this goal memory can be used to guide steering commands between different anchor points that are derived from it, using vector-based operations whose implementations are well understood in the fly central brain^35–37,71,72^. We do invoke a higher-dimensional goal memory than is thought to be implemented in the fly brain, and we assume that multiple distinct goal vectors can simultaneously be derived from it. Nevertheless, when taken together, these studies highlight the neural plausibility of our proposed algorithm, and underscore the power of vector-based computations in guiding flexible behavior.

### Outlook

The structure of behavior provides important clues for understanding how animals rapidly gather new information from their surroundings and modify their actions accordingly. Here, we provide insights into how this structured exploration can support active inference, hypothesis testing, and rapid learning in a manner that captures key aspects of behavioral data from mice learning a hidden target navigation task, efficiently navigating around novel obstacles, and adapting to new target locations. The underlying algorithms rely on computational elements that are shared across species, and that can be combined with additional map-like representations to support rapid learning and inference in more abstract task domains. Future work will be critical for producing refined predictions about neural representations that might be expected from such combined models, and for eventually testing those predictions in tasks that integrate learned trajectories with knowledge about the extrapersonal structure of the environment.

## Acknowledgements

We thank members of the Dudman and Hermundstad labs for helpful feedback on this work; Avni Garg for pilot analyses and simulations; Laura Grima, Luke Coddington, Brad Hulse, Ling-Qi Zhang, Wiktor Mlynarski, Albert Lee, and Vivek Jayaraman for feedback on the manuscript. WJ is supported by Shanghai Pujiang Program (23PJ1408600) and the National Natural Science Foundation of China (grant no. 32471088). JTD is a Senior Group Leader and AMH is a Group Leader at the Janelia Research Campus of the Howard Hughes Medical Institute.

## Author Contributions

JTD and AMH conceptualized the problem and conceived the framework. NYA wrote code to pilot the base computations, with guidance from JTD and AMH. WJ (Fig. 4, 5, 6), LTC (Fig. 1, 3), SG (Fig. 1, 3) collected all data reported here and wrote data loading code. JTD wrote all data analysis code and analyzed all data presented here, with input from AMH. AMH performed all derivations and wrote all simulation code used in the paper, with input from JTD. JTD and AMH wrote the paper, with input from all authors.

## Data Availability

All data and code used in analysis are available upon request. A curated dataset from which all figures were produced will be posted upon publication.

## Code Availability

All custom code for generating and simulating learning agents is available at https://github.com/HermundstadLab/trajectoryLearning. A secondary implementation that was used to pilot some of the base computations can be found at https://github.com/NadaYehia/Model-for-rapid-spatial-learning-via-efficient-exploration-and-structured-navigation

**Supplemental Figure 1:**
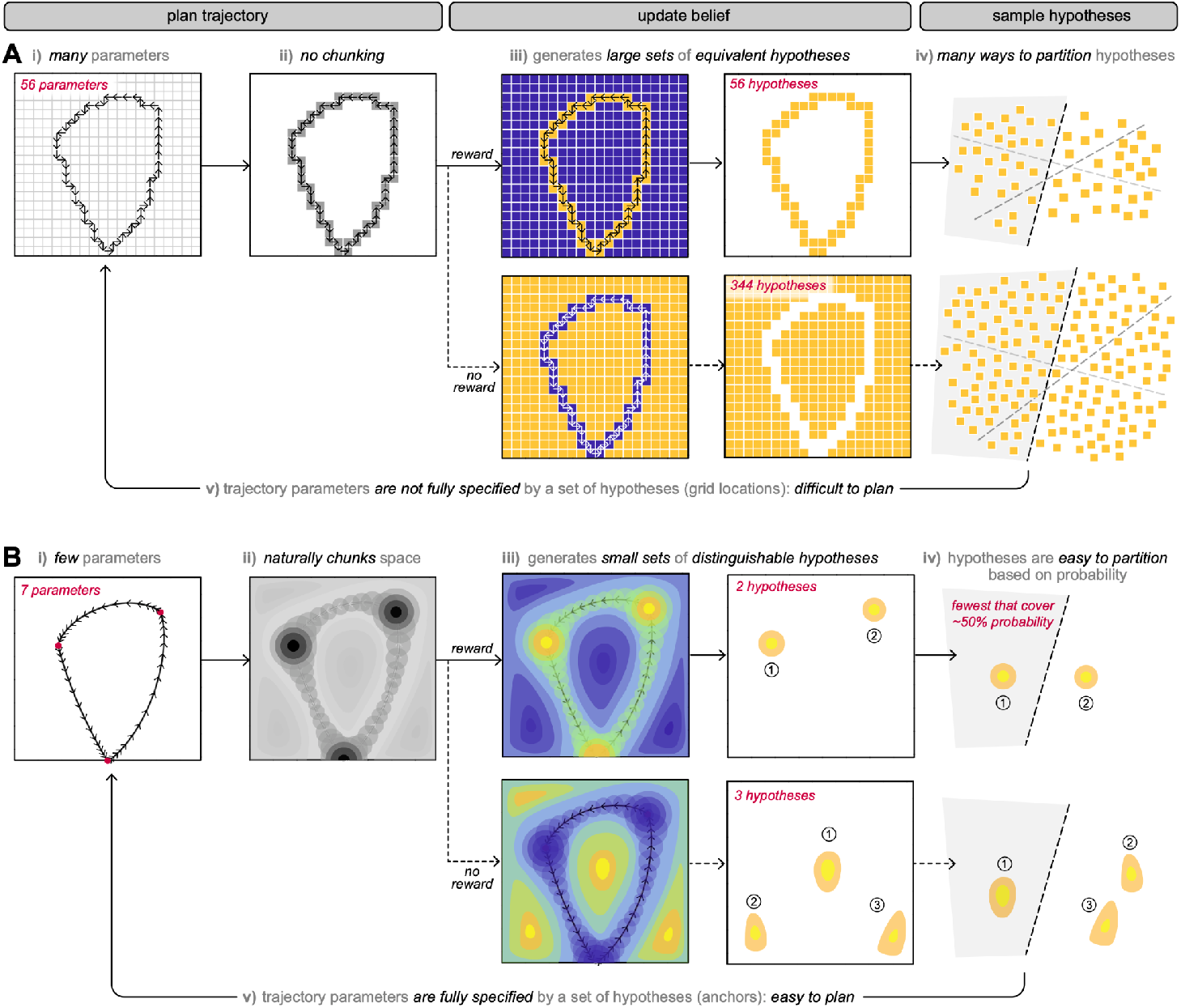
Tabular strategies suffer from multiple issues that slow learning. **A-B)** Comparison of a tabular representation (A) and a continuous representation (B) that share the same base modules introduced in the main text. For visualization purposes, both strategies are represented in cartesian coordinates, but can equivalently be transformed in polar coordinates, as shown in the main text. **A) i**. In a tabular representation, agents traverse space by taking one of an elemental set of actions to transition between grid locations. Generating a trajectory requires specifying a large set of elemental actions. i**i**. The resulting trajectory does not have any local temporal structure, and thus does not naturally chunk space. **iii**. As a result, if this trajectory is used to update a belief, it generates large sets of equivalent hypotheses based on whether it did (upper row) or did not (lower row) receive a reward. **iv**. Because the generated hypotheses are all equivalent, there are many ways to partition them into subsets to evaluate with subsequent trajectories. **v**. The subselected hypotheses take the form of specific grid locations; the trajectory planner must then plan a route between them. For a given set of grid locations, there are many different ways to plan a trajectory between them; in other words, a discrete and discontinuous set of grid locations does not fully specify the parameters of a trajectory that intercepts them, and planning is not straightforward. **B) i**. We chose to represent trajectories with a parametrized set of generative functions that use a small number of parameters to specify a continuous trajectory through space. **ii**. The resulting trajectory has both spatial and temporal structure, and naturally chunks space. **iii**. As a result, if this trajectory is used to update a belief, the resulting probability mass is locally higher or lower in regions where the agent spent more time. This results in a small and separable set of hypotheses given by the peaks of the posterior belief. **iv**. The height of these peaks reflects the time spent at different locations that ultimately led to reward; as a result, the peaks can be ranked by their posterior probability and easily partitioned. It is straightforward to select the smallest set of hypotheses that maximizes the expected information gain across both possible outcomes. **v**. The subselected hypotheses take the form of anchor points, which—together with an initial heading direction—fully specify the trajectory that intercepts them. As a result, planning is straightforward.

## Methods

Male and female mice, typically aged 3-9 months were used in this study. Mice were housed in a reversed 12:12 light/dark cycle (lights on at 18:00) and tested in the dark phase. All procedures were approved by the Janelia Research Campus Institutional Animal Care and Use Committee (IACUC) and were consistent with the standards of the Association for Assessment and Accreditation of Laboratory Animal Care.

### Behavioral Data

#### Hidden target task

Real-time position data from online tracked body centroid was recorded at approximately 50 Hz and was used either raw or synced with either a 100 Hz or 17 Hz clock on external recording equipment. The key hardware and custom software are described at http://dudmanlab.org/html/resources.html. Hardware was controlled with custom scripts written in the free software Processing (processing.org) and Arduino IDE (http://arduino.cc); data were analyzed with Matlab (http://mathworks.com). Occasional tracking errors were removed and interpolated over and then x,y position data was smoothed with a Savitzky-Golay filter. Position is reported relative to a reward located at *X* = 0, *Y* = 0. Video resolution was ~1 pixel/mm. The square arena was 75 cm per side.

To extract trajectories, we used a custom algorithm that used a threshold amplitude (7 cm) and minimum duration (~ 0.5 sec) to extract trajectories and find approximate start and stop frames of individual trajectories that originated near the reward port and returned to the reward port. Scalar statistics of each trajectory were then computed from the positions (or derived values, angle relative to reward port, velocity, distance, etc.) between event starts and stops.

#### Hidden target task and training

Mice were not pre-trained or shaped, rather task parameters remained constant from the first session in which naive mice were placed within the 75 × 75 cm hidden location arena. Sweetened water rewards (0.05 mg/mL Saccharin, Sigma) were delivered via a reward port (1.5 mm diameter metal tube) positioned 2 inches high at the center of one arena wall. A small plastic well was attached to the end of the tube to ensure water remained pooled near the tip of the tube. Reward delivery via the opening of a solenoid along the water line was triggered upon detection of the mouse within an unmarked 14 × 18 cm “target area” near the center of the arena. Mice then had to return to the reward port to collect reward, which also reset the task to the target-entry detection phase (this ensured that multiple rewards could not be triggered without returning to the reward collection area). Mice experienced one session per day, which was considered complete once the animal had collected ~1 mL of cumulative water reward. Some sessions early in training were terminated before the ~1 mL cutoff if 3 hours had passed. The reward delivery was controlled by a silent (in a subset of animals) or insulated solenoid valve outside the enclosure. In a subset of experiments, lick detection at the reward port was accomplished with a home-built capacitance detector and recorded synchronously at video rate to confirm accurate timing of reward collection.

To control execution of the task, a mouse’s position was recorded via a USB camera mounted below the clear floor of the enclosure. A real-time tracking algorithm was developed in which the video frame was converted to black and white, subtracting a blank background without a mouse, blurred, and then a standard OpenCV blob detection algorithm was applied with user customizable threshold settings, as described previously^5^. The center of the mouse body was calculated at every frame from the center of the detected blob, and a running buffer of positions were tracked by custom software written in Processing (www.processing.org) and written to a csv file. These position data were used for subsequent analysis of behavior. Reward delivery, real-time lick detection, and laser stimulation pulses were controlled with custom software using Arduino Mega hardware (www.arduino.cc). Sync pulses generated by the Arduino hardware were recorded in both behavior and parallel high speed recording systems to confirm robust behavioral timing and to synchronize data streams offline.

### Behavioral Data Analysis

Occasional tracking errors were interpolated over, and then position data were smoothed with a 3rd order Savitzky–Golay filter (MATLAB function “sgolayfilt()”). Following synchronization, behavior data were downsampled to 100 Hz or 17Hz for storage and analysis. Individual trajectories were defined by the mouse leaving a ~10 cm radius “reward collection area” around the reward port, and then re-entering following a minimum time threshold (0.3 s). Thus successful vs. unsuccessful trajectories were defined by whether reward was triggered between leaving and re-entering the reward collection area. Comparisons of trajectory features between different reward-size groups were made using matched numbers of trajectories from the beginning of each given session in order to compare animals at equivalent within-session states.

Custom code was written to extract reported parameters such as anchor points, posterior entropy, path length, and reward rate, similar to that described previously^5^. Anchor points were estimated by detecting peaks in the speed vs. time profile for each trajectory. If minima between peaks fell below a speed threshold, the first location of the minima was defined as an anchor. Thus, there are *N*_peaks_ *-* 1 anchors in a given trajectory. To estimate likelihoods from path trajectories, all sampled *x, y* positions were binned into 2D histograms on a 100 × 100 grid (~8~8 mm elements), smoothed with a gaussian kernel *(σ* =16mm), adjusted for reward/no reward, and normalized. Posterior belief estimates were built up in the manner described for the model implementation and normalized to the entropy of a flat posterior over the same grid.

## Computational Methods

Below, we provide a summary of key model assumptions. A full description of methodology can be found in the Supplemental Methods. All code was written in MATLAB 2023b (http://mathworks.com).

### Agent architecture

We constructed an agent that consists of three core modules for (1) maintaining and updating internal beliefs about the consequences of past trajectories, (2) using these beliefs to select anchor points for structuring future trajectories (and, when necessary, augmenting these anchor points to avoid obstacles), and (3) planning trajectories through the set of selected anchor points. We assume that the agent maintains its internal beliefs and selected anchor points in polar coordinates; these, in turn, are used to control its forward speed and angular velocity over time. We assume that the agent navigates with an arena of finite size, and that the agent itself occupies a finite width 6_*a*_. We use these two spatial scales to bound the agent’s internal belief and executed trajectories.

Below, we detail the construction of each of the agent’s core modules.

### Planning and Executing Trajectories

#### Parameterizing trajectories

Our agent controls its speed *v*(*t*) and angular velocity *ω*(*t*) over time *t* to travel between pairs of anchor points *{*(*r, θ*)*}* (note that these anchor points are specified in polar coordinates; see Section ‘Selecting Anchor Points’ below). We assume that the speed and angular velocity follow parameterized functions that are structured in time: *v*(*t*) = *A*(1 − cos(2*πt/T*))*/*2 and *ω*(*t*) = *m* (note that the constant angular velocity *m* leads to linear changes in the agent’s heading *ϕ*(*t*) = *ϕ*_0_ + *mt*. Together, these functions are fully specified by three parameters: *A, m, T*. The amplitude *A/*2 specifies the maximum speed along the trajectory segment, and *T* specifies the duration of the trajectory segment. We can thus choose parameter values that guarantee that the agent will begin and end its trajectory segment on a specified pair of anchors.

We make several simplifications to reduce the number parameters. (1) We assume that the duration of the trajectory segment scales with the vector distance *R* between two anchor points, such that *T* = *R/ρ*. (2) We assume that the trajectory is symmetric about the vector that connects two anchor points, such that heading as the agent leaves one anchor point (which we measure relative to this vector and denote *δ*) is the negative value of its heading upon arriving at the second anchor. (3) We assume that the heading is continuous across anchor points. Together, this means that we have to specify 2*N* − 1 parameters to generate a trajectory between *N* anchor points: the maximum speed and duration along each of *N* − 1 segments, and the initial heading angle at the start of the trajectory. We use the coordinates of the anchor points to constrain the maximum speed and duration of all segments (see SI Methods), and we optimize the initial heading to minimize the total path length along the entire trajectory (see *Optimizing trajectories to minimize path length*).

#### Ordering anchor points

To order anchor points, we solve a simplified version of the traveling salesman problem (alternatively, anchor points can be ordered by their angular orientation, which is simple, fast, and does not impact our overall conclusions). We apply this ordering to the *N*_*s*_ anchor points that are selected by the sampler module (see *Selecting anchor points*), and we then append the home port as an anchor point to the beginning and end of this ordered set. This gives a total of *N* = *N*_*s*_ +2 anchor points.

Prior to ordering the *N*_*s*_ anchor points, we sort them in either ascending or descending order (this is randomized on each trial to prevent the agent from traveling in a stereotyped direction). If *N*_*s*_ *<*= 5, we enumerate all possible orderings of these anchor points. We append the home anchor to the beginning and end of each ordering, compute the summed vector distance between successive pairs of anchor points, and select the ordering that minimizes this distance. As *N*_*s*_ increases, it becomes increasingly computationally costly to enumerate all sets of orderings. In cases where *N*_*s*_ *>* 5, and we break the full set of anchor points into *N*_*s*_*/*5 subsets, and we solve the traveling salesman problem within each subset. To this end, we find the best ordering of the first subset of anchor points. Then for each subsequent set of anchor points, we evaluate the summed vector distance by appending each possible ordering within the current to the best possible orderings of all previous sets.

#### Optimizing trajectories to minimize path length

The constraints described above—and in particular, enforcing that the heading is continuous across anchor points—can force the agent to run circuitous trajectories with long path lengths. To mitigate this, we allow the agent to execute its trajectory within some tolerance of each anchor point, if that allows the agent to run a shorter path. We do this by optimizing the placement of the anchors points in order to minimize the curvilinear path length along the trajectory. We constrain the final anchor points to be within some tolerance of the original anchor points; we set this tolerance to be a fraction *f*_tol_ of the maximum angular and radial span of the arena, where *f*_tol_ is set for each anchor individual depending on the width of the peak in the prior belief that was used to sample that given anchor point. For each peak within the belief distribution, we measure its width *σ*_peak_; we then measure *f*_tol_ = min(*σ*_max_, 0.05*σ*_peak_) (i.e., we compute 5% of the standard deviation of the peak, up to a maximum tolerance *σ*_max_). We then use this to set the angular and radial tolerance about each anchor point, with *r*_tol_ = *f*_tol_*r*_max_ and *θ*_tol_ = *f*_tol_*θ*_max_, where *r*_max_ and *θ*_max_ are the maximum radial and angular bounds of the arena measured relative to the home port. Each anchor point (*r*_*i*_, *θ*_*i*_) is then optimized with the bounds *r*_*i*_ *∈* [*r*_*i*,0_ − *r*_tol_, *r*_*i*,0_ + *r*_tol_] and *θ*_*i*_ *∈* [*θ*_*i*,0_ − *θ*_tol_, *θ*_*i*,0_ + *θ*_tol_], where (*r*_*i*,0_, *θ*_*i*,0_) are the initial coordinates of the anchor point.

#### Planning boundary trajectories

If a majority of the *N*_*s*_ anchor points lie along the boundary, the agent plans a boundary trajectory, rather than attempting to plan a continuous trajectory through a large set of boundary anchors. We define boundary anchors as those anchors within a tolerance Δ_*a*_*/*2 of the arena boundary, where Δ_*a*_ is the width of the agent (see *Agent architecture*). If more than *N*_*s*_*/*2 anchors are boundary anchors, the agent uses these boundary anchors (together with anchors at the home port and at the corners of the arena) to generate a boundary trajectory.

#### Evaluating trajectories

After planning a trajectory, we assume that the agent evaluates the quality of this trajectory against its internal error map (see *Updating error map*). This error map takes on negative values in places where planned trajectories were unsuccessful. If the minimum predicted error along the trajectory was less than a (negative) fixed threshold *E*_min_, the error map was used to select a set of anchor points to augment the trajectory. This criterion ensures that the agent only used this error map if the history of past trajectories had led to sufficiently large deviations between planned and executed trajectories. In these cases, the augmented anchor points were appended to the existing set that was sampled from the belief, and the agent re-planned (and re-optimized) a trajectory using the augmented set of anchor points. This second planning and optimizing step was only done once, and the result trajectory was then executed (see *Executing trajectories*).

#### Executing trajectories

We analytically computed the coordinates of planned trajectories, rather than numerically integrating the speed and heading profiles; as a result, they can be evaluated at arbitrary time discretizations. We used a finite discretization Δ*t*, such that there are *ρ/*Δ*t* time points per unit distance (where *ρ* is a scale factor; see *Parameterizing trajectories*). When executing planned trajectories that exceeded the arena bounds, we assume that any points that lie outside these bounds are mapped to the nearest boundary (accounting for the additional half width of the agent Δ_*a*_*/*2). When executing trajectories that encountered an obstacle, we diverted the agent’s trajectory around the obstacle until the agent could reconnect with previously planned anchor points. Upon first encountering the obstacle, we assumed that the agent turned in the direction (clockwise versus counterclockwise) that minimized its change in heading. To divert the trajectory around the obstacle, we considered three different conditions depending on whether the next planned anchor point was “ reachable” from a given face of the obstacle (i.e., whether there was an unobstructed path from all points along the obstacle face to the anchor point). (1) If the next anchor point was reachable from the face where the agent first encountered the obstacle, we assumed that agent traveled along the obstacle face until it encountered its originally-planned trajectory. In this case, we assumed that the agent used the corresponding entry and exit points as anchors to generate a boundary segment (see *Planning boundary trajectories*), before continuing its planned trajectory. (2) If the next planned anchor was not reachable from the face where the agent first encountered the obstacle, we assumed that the agent traveled along the obstacle until that anchor point first became reachable (which, for a rectangular obstacle, would be at a corner point). For this portion of the trajectory, we assumed that the agent used its entry point, together with the obstacle corners, as anchor points to plan a boundary trajectory. At this corner point where the agent exited the obstacle face, we assumed that the agent used the corner point and the next planned anchor point as two anchors for generating a trajectory segment; we assumed that the agent either made a directed run to the planned anchor (in which case the initial heading was *δ* = 0), or it peeled away from the obstacle boundary with an initial heading that was tangent to it. Between these two scenarios, we assumed the agent selected the one that minimized the change in heading upon encountering the next planned anchor point. (3) If the next anchor was not reachable because it fell inside the obstacle, we assumed that the agent followed the same procedure described above to travel to the subsequent planned anchor point.

As with the arena boundaries, we accounted for the agent’s finite width along obstacle boundaries, and assumed that the agent traveled along the boundary but at a distance Δ_*a*_*/*2 away from it.

#### Parameter settings

For all results shown in the main text, we used the following parameter settings: *ρ* = *r*_max_*/*2; *σ*_max_ = 0.005; *E*_min_ = −0.05; Δ*t* = 0.01. These parameters are specified in the agent’s ‘planner’ data structure.

### Maintaining and Updating Internal Beliefs

#### Updating a belief about the successful anchor

Our agent maintains an internal belief about a candidate point that will generate a successful trajectory. The agent represents this anchor as a polar vector defined relative to a fixed reference point; for all results shown in the main text, we assumed that the reference point was the home port. We assume that the agent knows the size of its environment; for closed environments like the arena considered here, this can be inferred by running a boundary trajectory around the environment. The agent then maintains a belief over possible anchor vectors within the environment, defined in polar coordinates as a function of the angle *θ* and distance *r* measured relative to the home port. We refer to this as the “anchor belief”, to distinguish it from a belief about the current environmental context.

The agent iteratively updates this anchor belief over time based on all past trajectories 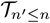 and outcomes 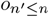 (where *n* indexes the current trial, and *n*^*′*^ ≤ *n* denotes the history of past trials up to and including the current one). After running a trajectory 𝒯_*n*_ through a set of vectors *{*(*r, θ*)*}*_*n*_ and observing an outcome *o*_*n*_, the agent computes the likelihood of observing the current outcome under different possible anchor placements, and uses this likelihood to update its belief 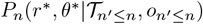 about the successful anchor vector (*r*^*∗*^, *θ*^*∗*^). This construction assumes that the agent knows that there is a single anchor point that can be used to reliably generate a rewarding trajectory. We also assume that the agent believes that the probability of receiving a reward falls off with distance from the correct anchor placement. Thus, if the agent executes a rewarding trajectory, the agent knows that the successful anchor is likely to be along the executed path, and unlikely to be far away. We construct this likelihood *L*(*o*_*n*_) = *P* (*o*_*n*_|*r*^*∗*^, *θ*^*∗*^, 𝒯_*n*_) by convolving a Gaussian kernel with the executed trajectory; we set the width *σ*_*L*_ of this Gaussian such that relative changes in angular and radial coordinates are given equal resolution. This construction assumes that the agent can track its path-integrated orientation and distance relative to home. We then scale the resulting function 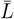 to lie between ℓ and 1 − *f*. The likelihood of a rewarding outcome is then 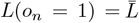, and the likelihood of an unrewarding outcome is 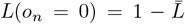. This likelihood is then multiplicatively combined with the prior belief and normalized to obtain the updated posterior belief: 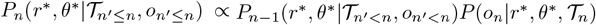. After a rewarded trial, this update results in localized probability mass around executed anchor points where the agent spent more time (and conversely, after an unrewarded trial, this pushes probability mass away from executed anchor points).

We defined the anchor belief over the allowed set of anchor vectors within the arena, such that *θ ∈* [0, *π*] and 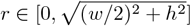, and where *w* and *h* are the width and height of the arena, respectively. We discretized *r* and *θ* into *n* evenly-sized bins (such that *n*_*r*_ = *n*_*θ*_ = *n*, and such that the full ranges of each axis were evenly tiled), and we defined our belief as a grid of (*n*_*θ*_ *× n*_*r*_) points. We accounted for the agent’s finite width Δ_*a*_ by masking those points that fell beyond a tolerance Δ_*a*_*/*2 of the arena bounds (i.e., we masked any points that satisfied *x < x*_min_ + Δ_*a*_*/*2, *y < y*_min_ + Δ_*a*_*/*2, *x> x*_max_ − Δ_*a*_*/*2, or *y > y*_max_ − Δ_*a*_*/*2).

#### Updating a belief about the underlying context

In addition to maintaining a belief about the successful anchor vector, we assume that the agent maintains an internal belief about the current context. Individual contexts correspond to anchor beliefs that have been previously stored in a memory cache (see *Using surprise to cache the existing anchor belief and initialize a new prior*); thus, the context belief is a vector of probabilities—one for each context in the cache—that sum to one. To update the context belief, the agent computes the probability that the executed trajectory would have led to the observed outcome *o*_*n*_ under each anchor belief in the cache. For a given prior belief 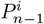 about the *i*^*th*^ anchor vector (which corresponds to the *i*^*th*^ context *C*_*i*_), the outcome probability is given by: 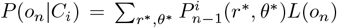 (note that this is equivalent to the normalization factor in the Bayesian update). This outcome probability is then combined with the prior context belief to obtain the updated posterior belief: *P*_*n*_(*C*_*i*_) ∝ *P* (*o*_*n*_|*C*_*i*_)*P*_*n*−1_(*C*_*i*_) (see Supplemental Methods for more details).

#### Using surprise to cache the existing anchor belief and initialize a new prior

The agent can use its prior anchor belief and executed trajectory to evaluate the surprise of the outcome it received. We compute this surprise as *S*(*o*_*n*_|*P*_*n*_) = − log(*P* (*o*_*n*_)), where *P* (*o*_*n*_) is the probability under the current prior that the executed trajectory led to the observed outcome, as described above in *Updating a belief about the underlying context*. We evaluate this surprise relative to the surprise of the same outcome under a uniform prior, *S*(*o*_*n*_|*P*_0_). If the relative surprise 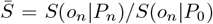 exceeds a threshold 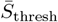 (i.e., if the outcome is sufficiently more surprising under the current prior compared to a uniform prior), the agent initializes a new (uniform) prior belief. To prevent unnecessary resets (i.e., in cases where the anchor belief is localized in a manner that generates consistently rewarding trajectories, and the executed trajectory falls off of the target for a single trial), we assume the agent use the history of surprise values on the past Δ*n* trials to detect a target switch.

When the agent initializes a new anchor belief, we assume that it can preserve its prior anchor belief in a memory cache. Under this formulation, each new initialization of the prior will increase the size of the cache, and thus the number of possible contexts with which the agent must contend. When initializing the anchor belief, the agent must also initialize the corresponding context belief. We assume that in initializing a new anchor belief (and thus a new context), the agent believes that this context is as or more likely than previous contexts; it thus assigns a probability of 0.5 to the newly initialized context, and renormalizes the existing context prior to make up the remaining 0.5 probability.

We assume that the agent has finite memory, and can only retain a maximum cache size of *N*_c,max_. In scenarios where the agent is already occupying all of its available cache entries and decides to cache its existing belief, we assume that it begin over-writing an existing anchor belief. We assume that the agent selects the anchor belief that is closest to uniform (measured via the KL divergence between the anchor belief and a uniform prior). Because we assume that cache entries decay over time via a memory leak (see *Memory decay* below), this will typically have the effect of over-writing the entry that was updated most distantly in the past.

Finally, we assume that the agent can sample anchor points from previously cached beliefs (see *Selecting the appropriate prior belief from which to sample*) in order to generate trajectories, but we assume that the agent always uses the outcome of its trajectory to update its current belief (i.e., it preserves previously cached memories without altering them).

#### Updating an error map

We assume that the agent can track differences between planned and executed trajectories using the same path integration mechanisms that it uses to construct its likelihood function. When these two trajectories differ from one another (for example, when the agent encounters an obstacle), the agent uses these differences to track its error over time. The agent stores this information in an “error map” *E*(*r, θ*) that updates after each erroneous trial. We assume a simple update: *E*_*n*_(*r, θ*) = *E*_*n*−1_(*r, θ*) + *L*_exec_(*o*_*n*_ = 1) − *L*_plan_(*o*_*n*_ = 1), where *L*_exec_(*o*_*n*_ = 1) and *L*_plan_(*o*_*n*_ = 1) are the likelihood of a rewarding outcome under the executed versus planned trajectories, respectively. Thus, portions of the planned trajectory that could not be executed will result in negative error values, and portions of executed trajectory that enabled the successfully maneuver around obstacles will result in positive error values.

#### Parameter settings

For all results shown in the main text, we used the following parameter settings: *n* = 100, ℓ = 0.05, *σ*_*L*_ =.075*n* (in units of discrete elements of the belief distribution), 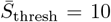 (i.e., the outcome is 10 times more surprising under the current prior compared to a uniform prior), Δ*n* = 2, *N*_c,max_ = 5. These parameters are specified in the agent’s ‘belief’ data structure. Parameters for the arena and agent size are discussed below. All simulations were initialized with a flat anchor belief.

### Selecting Anchor Points

#### Selecting the appropriate prior belief from which to sample

The agent uses a prior belief about the anchor vector to sample anchor points that will guide future trajectories. In cases where the agent does not allow for the possibility of multiple environmental contexts (i.e., when it does not maintain a memory cache), the agent always samples from its current anchor belief. However, in cases where the agent maintains a memory cache, we assume that the agent first samples a context from its memory cache, and then sample anchors from the anchor belief specified by that context. There are, in principle, several different ways to implement this sampling; for the results shown here, we assumed that the agent sampled the context in proportion to its prior probability, as stored in the context belief (see *Updating a belief about the underlying context*).

#### Selecting anchor points from the prior belief

Our agent uses peaks in its prior belief to select anchor points on a given trial. We identified these peaks using the MATLAB function *peaks2*.*m* created by Kristupas Tikuišis (available at Matlab Central File Exchange, file 113225). We selected those peaks that exceeded a specified minimum peak height *P*_min_ and minimum peak separation 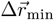. We normalized the set of peak heights to sum to one; this allowed us to use these peak heights as a proxy for the full belief space. We then rank-ordered the peaks by their heights, and selected the minimal set that, when summed, exceeded a probability of 0.5.

If the prior belief was sufficiently flat, we instead randomly sampled a fixed set of *N*_init_ anchor points. We first computed the KL divergence *D*_KL_(*P*_*n*_||*P*_0_) between the current prior *P*_*n*_ and the initially flat prior *P*_0_; if this was smaller than a fraction *f* of the base entropy *H*_0_ of the flat prior (i.e., *D*_KL_(*P*_*n*_||*P*_0_) *< fH*_0_), the current prior was deemed sufficiently flat, and the agent randomly sampled a set of anchor points.

This returns a set of *N*_*s*_ sampled anchor points 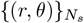, specified in polar coordinates.

#### Selecting anchor points from the error map

In cases where the agent used an internal error map to augment this set of anchor points (see *Updating error map*), we used the same sampling procedure described in the previous paragraph. However, rather than selecting anchor points that covered half of the peak probability distribution, the agent selected all peaks (i.e., covered the full peak probability distribution). To extract these peaks, we first transformed the error map into a probability distribution by removing non-negative elements and normalizing the resulting distribution to sum to one, and proceeded as described above.

#### Parameter settings

For all results shown in the main text, we used the following parameter settings: 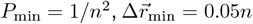 (in units of discrete elements of the belief distribution), *f* = 0.001, *N*_init_ = 8 anchor points. Note that we chose the initial number of anchors points such that half of the agents on average intercepted the target on the first run. These parameters are specified in the agent’s ‘sampler’ data structure.

### Simulations

#### Arena configuration

For all simulations shown in the main text, we constructed a square arena of size *w* = *h* = *H*, with *H* = 10. The home port was positioned at *x* = *y* = 0. All targets were of size *w*_*t*_ = *h*_*t*_ = 0.1*H*. For simulations involving a single target, we centered the target at *x*_*tc*_ = 0 and *y*_*tc*_ = *H/*2. For simulations involving 5 targets, we position each target at a radial distance of *H/*2 from the home port, with angular orientations of [*π/*2, *π/*4, 3*π/*4, 3*π/*8, 5*π/*8] (such that target centers were given by *x*_*tc*_ = [0, 0.35, −0.35, 0.19, −0.19]*H*, and *y*_*tc*_ = [0.5, 0.35, 0.35, 0.46, 0.46]*H*). For simulations involving obstacles, we used an obstacle of size *w*_*o*_ = 0.4*H*, and *h*_*o*_ = 0.04*H*, centered at *x*_*oc*_ = 0 and *y*_*oc*_ = 0.25*H*. We assumed that the agent occupied a finite width Δ_*a*_ = 0.05*H*.

#### Learning simulations

Unless otherwise stated, we simulated batches of 500 agents and averaged the results across agents. All agents began with a uniform belief distribution over candidate anchors. On a given trial, agents were rewarded if any portion of their trajectory intercepted the target. To compute the reward probability for an individual agent, we performed a running average of binary rewards, using the Matlab function *movmean*.*m* with a window size of 7 trials and centered on the current trial. We computed this separately for blocks of trials corresponding to the same target or obstacle condition; in other words, we did not smooth rewards across changes in the arena configuration. This running average was not applied to any of the other belief or trajectory measures, with the exception of the comparison in Fig. 1C, where we performed a running average of the prior entropy and the number of anchor points for individual agents, and displayed their relationship over the first 10 trials of learning. We computed the entropy of the belief distribution in bits: *H* = *P*_*n*_ log_2_ (*P*_*n*_), and normalized by the entropy of a flat prior, *H*_0_ = − *P*_0_ log_2_(*P*_0_).

## Supplemental Methods

We consider an agent that uses a parameterized behavioral policy to generate navigational trajectories. We use this agent to study learning in a hidden target navigation task in which the agent must intercept a localized target somewhere in an open arena in order to receive a reward at a home port. The agent does not receive any localizing feedback about the target, and thus must infer how to generate behavior that reliably leads to reward.

To navigate through the environment, we assume that the agent uses spatial anchor points to specify segments of a trajectory. Given a pair of anchor points, the agent determines the parameters of a controller that can intercept the pair of anchor points. The agent can then generate a full trajectory by composing a sequence of anchor points and the smooth path segments between them. Guiding behavior then consists of appropriately selecting anchor points, and appropriately guiding trajectory segments between them.

We assume that the goal of the agent is to generate a compact trajectory that reliably triggers reward. To guide the selection of anchor points, we assume that the agent maintains and updates an internal belief about a single anchor point that can be used to generate a successful trajectory. This internal belief is used to specify hypotheses about candidate anchor points that could be successful; a downstream sampler then selects which of these hypotheses (in the form of anchor points) will be evaluated on the trajectory. Finally, the trajectory planner uses these anchor points to generate the egocentric speed and heading of the agent over time. This executed trajectory and its outcome are used to update the belief.

In what follows, we first detail our parametrization of behavior, and derive a set of generative equations for controlling the agent’s speed and heading over time and thereby generating navigational trajectories. We then derive the ideal Bayesian observer that uses these trajectories, together with their outcomes, to infer the placement of a successful anchor point. To aid in the inference, we couple this ideal observer with a sampler that selects a set of candidate anchors from the observer’s belief distribution. Finally, we discuss how these anchors are used by a trajectory planner to control future behavior.

### Notation

We use *n* to index trials, *i* to index parameters of a trajectory on a given trial, and *t* to index time along a trajectory in a given trial.

### Parameterization of behavior

We consider an agent that navigates through space using parameterized navigational trajectories. We assume that the agent controls its angular velocity *ω*(*t*) and forward speed *v*(*t*) to generate a segment of a trajectory; multiple segments can then be composed together to generate a full trajectory. We assume that each segment is defined by an increase and decrease in forward speed and a constant angular velocity, executed over a time interval *T*; we parameterize these using the following generative functions:

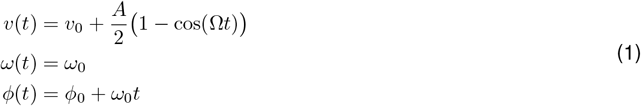

where *A/*2 *>* 0 specifies the maximum speed (measured relative to the initial speed, *v*_0_), Ω = 2*π/T* ensures that the speed will return to the initial speed at a time *T*, and *ω*_0_ specifies the constant angular velocity. Note that the constant angular velocity leads to a linear change in heading *ϕ* (*t*), as given above. The form of these generative equations is qualitatively similar to that used in Jiang, Xu, & Dudman 2022, with some modifications for flexibility and analytic tractability.

When integrated over time, Eq. produces arc-like trajectory segments that can be used to connect a pair of anchor points (*r*_*i*_, *θ*_*i*_) and (*r*_*i*+1_, *θ*_*i*+1_). Given a set of anchor points, we can then derive the set of control parameters *{v*_0*i*_, *A*_*i*_, *ϕ*_0*i*_, *ω*_0*i*_, *T*_*i*_*}* that will generate trajectory segments to connect them. For ease of notation, we will use *v*_*i*_(*t*), *ω*_*i*_(*t*), and *ϕ*(*t*) to denote Eq. 1 evaluated using the set of parameters *{v*_0*i*_, *A*_*i*_, *ϕ*_0*i*_, *ω*_0*i*_, *T*_*i*_*}*.

In what follows, we will outline the simplifying assumptions that use to constrain the set of trajectory parameters. We will use these assumptions to derive expressions for the maximum speed *A/*2 and the angular velocity *ω*_0_. We will then integrate the generative functions to derive the position of the agent over time, and show that these equations can be fully specified by a set of anchor points, together with the initial heading offset at the beginning of the trajectory. Finally, we will compute the curvilinear distance along the trajectory, which we will use to optimize the initial heading offset.

#### Simplifying assumptions

In practice, we will make several simplifying assumptions to constrain the set of parameters {*v*_0*i*_, *A*_*i*_, *ω*_0*i*_, *T*_*i*_}. For ease of notation, we assume that each trajectory segment *i* begins at time *t* = 0 and ends at time *t* = *T*_*i*_:

- We take *v*_0*i*_ = 0 ∀_*i*_, so that each trajectory segment is delineated by pauses at the anchor points.
- We assume that the initial and final heading directions are symmetric relative to the vector connecting two anchors. We will use Θ_*i*_ to denote the angle between two anchors, which is given by:

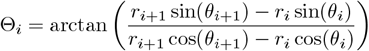 We use *δ* ∈[™*π, π*] to denote the heading direction relative to the vector connecting two anchors, such that *δ*_*i*_(*t*) = *ϕ*_*i*_(*t*) ™ Θ_*i*_. Requiring the initial and final headings to be symmetric about this vector means that *δ*_*i*_(*T*_*i*_) = ™*δ*_*i*_(0), or equivalently that *δ*_*i*_(*T*_*i*_) ™ *δ*_*i*_(0) = ™2*δ*_*i*_(0). Thus, the angular velocity can be written as *ω*_0*i*_ = (*ϕ*_*i*_(*T*_*i*_) ™ *ϕ*_*i*_(0))*/T*_*i*_ = (*δ*_*i*_(*T*_*i*_) ™ *δ*_*i*_(0))*/T*_*i*_ = ™2*δ*_0*i*_*/T*_*i*_, and the initial heading offset can be written as *ϕ*_0*i*_ = *δ*_0*i*_ + Θ_*i*_. Together, this gives *ϕ*_*i*_(*t*) = *ϕ*_0*i*_ + *ω*_0*i*_*t* = *δ*_0*i*_ + Θ_*i*_ *™* 2*δ*_0*i*_*t/T*_*i*_. In practice, we will go back and forth between *ϕ*_*i*_(*t*) and *δ*_*i*_(*t*) for mathematical convenience.
- We require that the agent’s speed and heading be continuous across pairs of anchors. With *v*_0_ = 0, the speed profile in Eq. already ensures that the speed is continuous. For the heading to be continuous, we choose *ϕ*_0*i*_ = ϕ_*i*−1_(*T*_*i*−1_), or equivalently *δ*_0*i*_ = *δ*_*i*−1_(*T*_*i*−1_) ™ (Θ_*i*_ *™* Θ_*i*−1_). For a set of anchors {(*r*_*i*_, *θ*_*i*_)}, this constraint means that we need only specify *δ*_00_ (i.e, the initial heading direction at the very start of the trajectory), and all other *δ*_0*i*_,*i >* 0, will be determined by *δ*_00_ and by the positions of the anchors.

With these assumptions, Eq. 1 becomes:

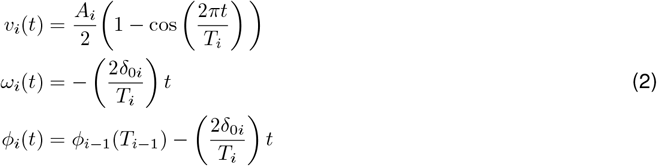

with *δ*_0*i*_ = *δ*_*i*−1_(*T*_*i*−1_) ™ (Θ_*i*_ *™* Θ_*i*−1_). With these assumptions, and given a set of anchor points, the trajectory is fully specified by the maximum speed *A*_*i*_ along each segment, the total time *T*_*i*_ taken to execute each segment, and the initial heading direction at the beginning of the trajectory, *δ*_00_. Thus, an *N ™*anchor trajectory generates *N ™* 1 trajectory segments, and is specified by 2*N ™* 1 parameters: *N ™* 1 parameters that specify the maximum speed along each segment, *N ™*1 parameters that specify the duration along each segment, and one parameter that specifies the initial heading offset.

#### Deriving the remaining parameters of the generative functions

We used the assumptions laid out in the previous section to derive the constant angular velocity *ω*_0*i*_. The maximum speed along the trajectory, which is specified by the amplitude *A*_*i*_*/*2, can be found by inverting the following equation:

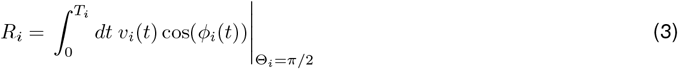

Note that this is the equation for *R*_*i*_ = Δ*x*_*i*_(*T*_*i*_) = ∫ *dt v*_*xi*_(*t*), with *v*_*xi*_(*t*) = *v*_*i*_(*t*) cos(ϕ_*i*_(*t*)), which is equivalent to the Euclidean distance between two anchors separated by an angle of Θ_*i*_ = *π/*2 rad. Solving this equation yields:

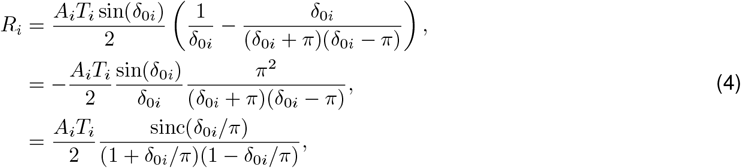

where we have used sinc(*x*) ≡ sin(*πx*)*/πx* (note that this is the implementation used in MATLAB). Thus, the maximum speed is given by:

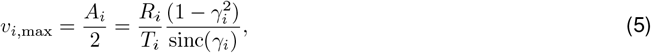

where we have defined *γ*_*i*_ = *δ*_0*i*_*/π*.

One can see from Eq. that the maximum speed increases with the distance between anchors and decreases with the time taken to travel between anchors. If we choose the timing function to be linear with distance, i.e., *T*_*i*_ = *f* (*R*_*i*_) = *R*_*i*_*/ρ*, then *A*_*i*_ depends only the initial heading direction *δ*_0*i*_ and the constant scaling factor *ρ*. Here, the constant *ρ* specifies the distance that can be covered per unit time. We will return to this assumption after we derive the final form of the trajectory equations.

#### Deriving the position of the agent over time

We can now compute the cartesian coordinates of the agent’s position over time by integrating the expressions given in Eq. 2. Note that this derivation is for our purposes only, and is not needed by the agent to generate and control its own behavior. Having closed-form expressions of the agent’s position allows us to simulate the behavior more efficiently while minimizing errors from numerical integration.

The cartesian components of the velocity are given by:

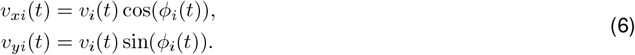

We can integrate these to get the cartesian components of the agent’s position over time:

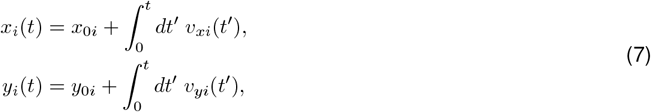

which gives:

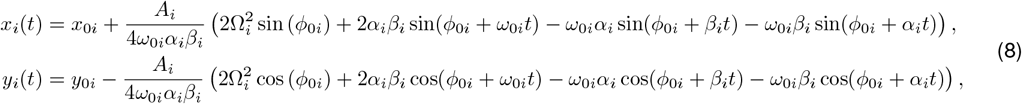

where we have defined the temporary variables *α*_*i*_ = *ω*_0*i*_ + Ω_*i*_ = Ω_*i*_(1 − *γ*_*i*_) and *β*_*i*_ = *ω*_0*i*_ − Ω_*i*_ = −Ω_*i*_(1 + *γ*_*i*_), and where *ω*_0*i*_ = −Ω_*i*_*γ*_*i*_ and Ω_*i*_ = 2*π/T*_*i*_.

As we did for the expression of the amplitude in Eq. 5, we will simplify these expressions to show that they remain finite as *δ*_0_ → *π/*2. First, we can rewrite Eq. 9 as follows:

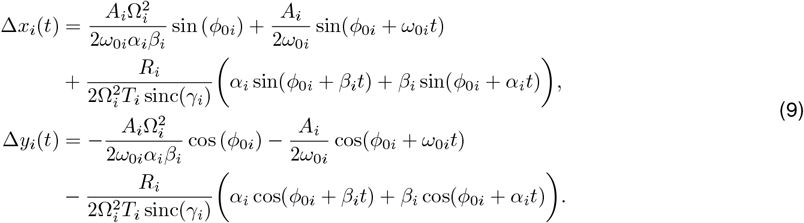

From this, one can see that the second set of terms in each expression is finite as *δ*_0_ → *π/*2. We can simplify the first two terms as follows (we will focus first on the expression for Δ*x*_*i*_(*t*)):

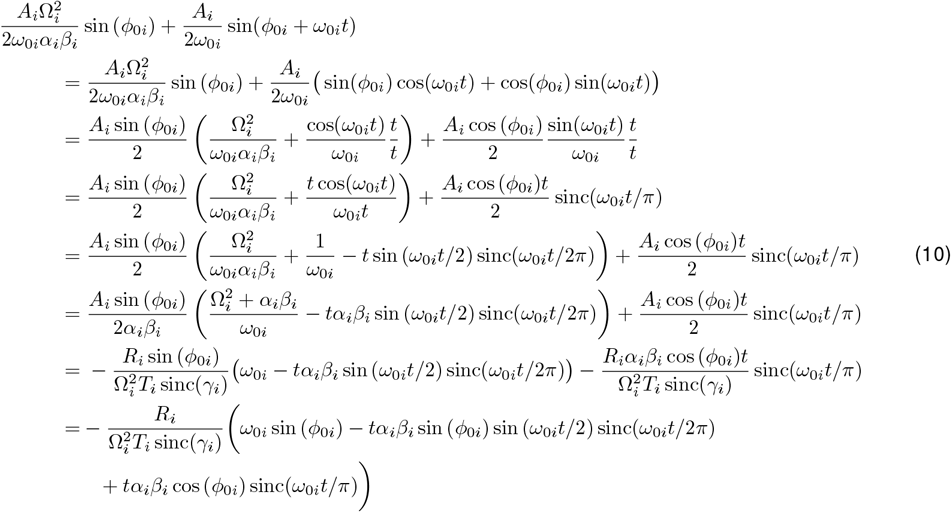

where we have used:

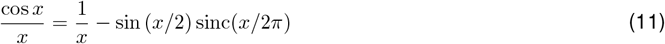

We can now write the full expression for Δ*x*_*i*_(*t*):

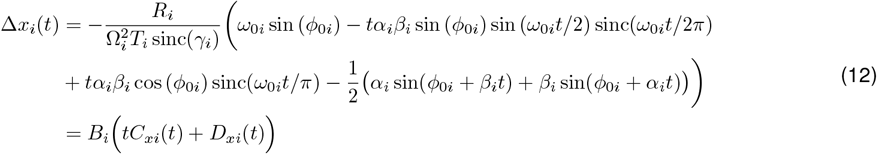

where we have made the substitutions *R*_*i*_*/T*_*i*_ = *ρ, ω*_0*i*_ = −Ω_*i*_*γ*_*i*_, *α* = Ω_*i*_(1 − *γ*_*i*_), and *β* = −Ω_*i*_(1 + *γ*_*i*_), such that we can write:

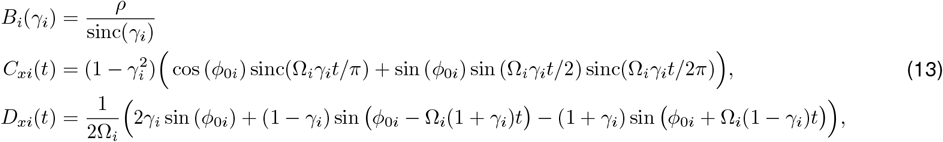

We can repeat the same calculations for the first two terms in the expression for Δ*y*_*i*_(*t*) in Eq. 9:

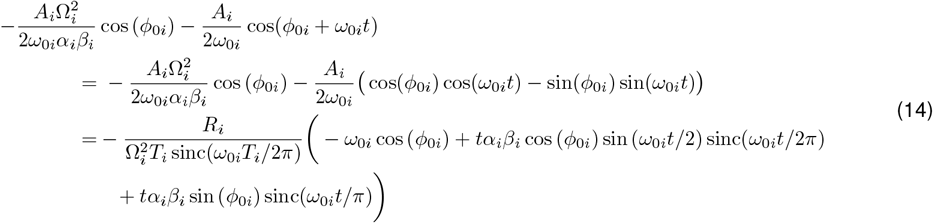

where we have used the observation that if we take cos(*∫*_0*i*_) → ™ sin(*∫*_0*i*_) and sin(*∫*_0*i*_) → cos(*∫*_0*i*_) in the second line above, we recover the second line of Eq. 14.

The full expression for Δ*y*_*i*_(*t*) is then:

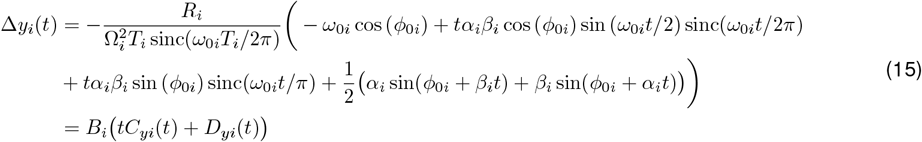

where we can again write

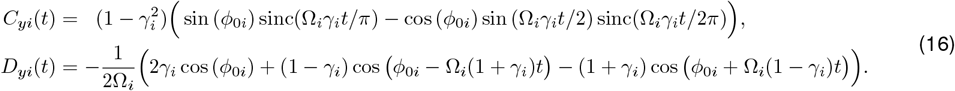

#### Full trajectory equations

Putting everything together, this means that we can fully specify a trajectory 𝒯(*t*;{(*r*_*i*_, *θ*_*i*_)}) given a set of anchor points {(*r*_*i*_, *θ*_*i*_)}. The instantaneous speed *v*_*i*_(*t*), angular velocity *ω*_*i*_(*t*), heading *ϕ*_*i*_(*t*), and position (*x*_*i*_(*t*), *y*_*i*_(*t*)) along each segment *i* of the trajectory can be written in closed form as follows:

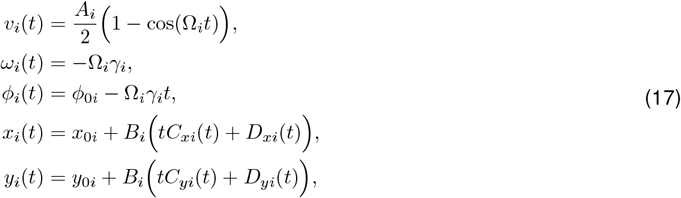

where

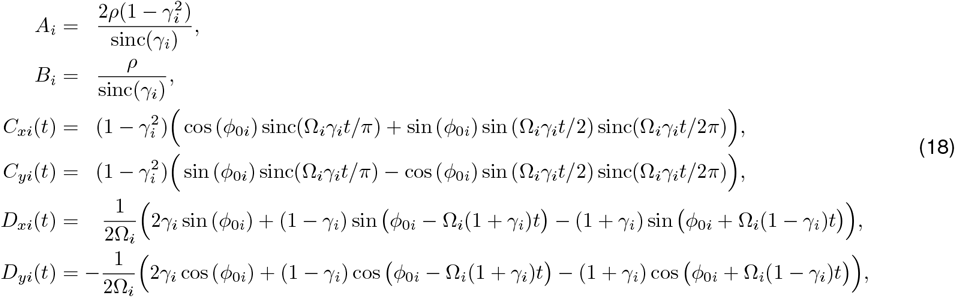

and where

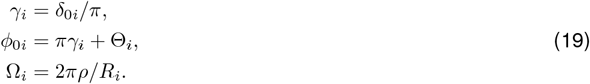

The initial conditions are given by:

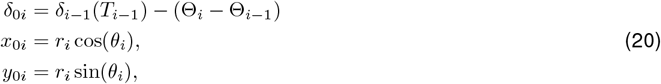

and the separation between anchors is given by:

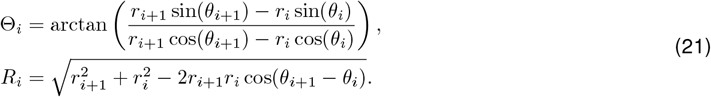

From these equations, it is easy to see that an *N ™* anchor trajectory is fully specified by the *N ™* 1, two-dimensional vectors between the anchor points {(*r*_*i*_, *θ*_*i*_)}, and the single initial heading angle *δ*_00_ at the beginning of the trajectory. Without additional constraints, we are free to choose *δ*_00_. In practice, for a given set of anchors {(*r*_*i*_, *θ*_*i*_)}, different choices of *δ*_00_ will lead to more or less efficient trajectories through those anchors. We therefore choose *δ*_00_ to minimize the curvilinear distance along the trajectory (i.e., path length) that passes through {(*r*_*i*_, *θ*_*i*_)}. More generally, we can use this idea to balance the efficiency and accuracy of a given trajectory by allowing the trajectory to pass within a given radius *r*_tol_ about each anchor. Increasing *r*_tol_ can make it possible to find more efficient trajectories at the cost of deviating from the chosen anchor points. We use this approach to simultaneously find a set of approximate anchors 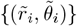 and an initial heading direction *δ*_00_ that minimizes path length for a given *r*_tol_.

#### Path length along trajectory

As described above, we can choose the initial heading angle *δ*_00_ to minimize the path length along the entire trajectory. For a given trajectory segment *i*, the path length can be computed analytically:

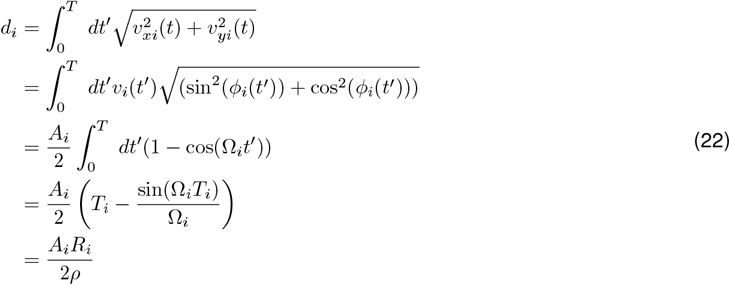

We see that under our assumptions, the path length along a trajectory segment *i* is a simple scaled product of the maximum speed *A*_*i*_ along the segment, and the distance *R*_*i*_ between the two anchor points connected by the segment. Different choices of *δ*_00_ will impact the maximum speed along different segments, and thus the total length *d*; by adjusting *δ*_00_, we can find the trajectory that minimizes path length while still intercepting the required set of anchors *{r*_*i*_, *θ*_*i*_*}*.

### Ideal Bayesian observer

We derive an ideal Bayesian observer that uses the history of past trajectories 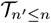 and outcomes 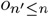 to infer the candidate anchor point 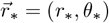 that generates a rewarding trajectory. This formulation assumes that the agent knows that a single anchor point is sufficient to generate a rewarding trajectory, but does not know where the anchor point should be placed. Instead, it must use the outcomes of past actions to infer the best anchor placement.

On a given trial *n*, we assume that a trajectory 𝒯_*n*_ causes the agent to pass through a set of points 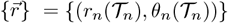 that are a radial distance *r*_*n*_ and angular distance *θ*_*n*_ away from the home port. This trajectory either leads to reward (*o*_*n*_ = 1) or no reward (*o*_*n*_ = 0), which can be used in combination with the executed trajectory _*n*_ to update the posterior belief 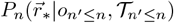. As we will discuss in the next section, we will use the posterior belief to select anchors points that specify the trajectory to be taken on the next trial; for now, we don’t make any assumptions about how the trajectory is generated.

For a given history of trajectories 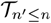, the agent’s posterior belief can be written as:

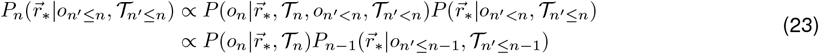

where we have used the fact that the outcome delivered on trial *n* depends only the executed trajectory 𝒯_*n*_ and the correct anchor 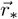, and does not depend on previous trajectories or outcomes. This allows us write the likelihood as 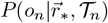. Similarly, the prior does not depend on the current trajectory 𝒯_*n*_, and can thus be written as the posterior 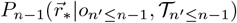 from the previous trial.

As a simple starting point, we consider a trajectory that consists of a single point 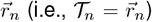, and assume a Gaussian likelihood centered about 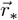:

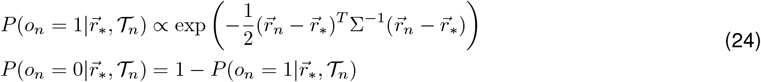

where

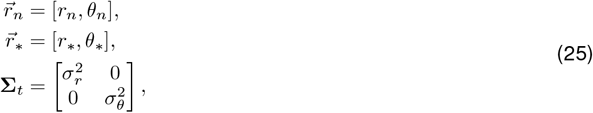

and where the proportionality denotes that we need a scaling factor to keep the likelihood bounded between 0 and 1. This likelihood captures the knowledge that the probability of receiving a reward is maximal (and equal to 1) when the agent passes through the point 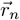 that aligns with the optimal anchor placement 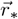, and falls of as a function distance between them. If, instead of passing through a single point, a trajectory passes through a set of points 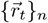, the likelihood becomes a sum of Gaussians centered on these points (equivalent to convolving a Gaussian function with the trajectory 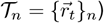:

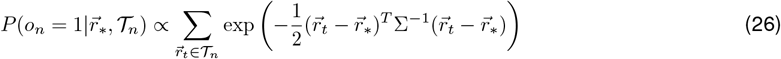

where *t* indexes points along a trajectory, as in the generative model above. This assumes that the agent can track its path-integrated orientation and distance over time as it executes a trajectory. Note that depending on the spacing of points along the trajectory *{r*_*t*_*}*_*n*_, this likelihood function can be multi-peaked.

When the belief update specified by Eq. 23 is iterated over time, the posterior will converge on an anchor placement that, when used to generate a trajectory, will reliably yield reward. Before convergence, the posterior represents the set of hypothesized anchors that the agent believes will generate reward. Next, we discuss how this information can be used to generate appropriate trajectories in order to evaluate these hypothesized anchors.

### Sampler

We can now combine the ideal Bayesian observer with the generative model to guide future trajectories. On a given trial *n*, we assume that the posterior *P*_*n*_(*r*_*∗*_, *θ*_*∗*_) can be used to select a set of anchors *{*(*r*_*i*_, *θ*_*i*_)*}*_*n*_, which can in turn be used to specify the trajectory 𝒯(*t*; *{*(*r*_*i*_, *θ*_*i*_)*}*_*n*_) that intercepts these anchors.

On a given trial, the posterior belief represents the agent’s knowledge about the successful anchor point. If the goal of the agent is to localize this posterior as quickly as possible (as we assume here), the optimal strategy is to cut the posterior probability in half on every trial. In the current setting, this requires executing a trajectory that passes through half of the probability mass of the posterior. As an approximation to this optimal case, our agent selects actions based on the peaks of the posterior, rather than on the full distribution. Each peak can be viewed as representing a distinct hypothesis about the successful anchor. The stronger the peak, the more confident the agent is about the corresponding hypothesis.

In such scenarios, if there are *N* candidate hypotheses captured by *N* distinct peaks in the posterior, the fastest way to narrow down these hypotheses is to select an action that eliminates half of the probability mass captured by these peaks. This can be achieved by using a subset of of these peaks as anchor points, which are then are used to plan and execute a trajectory through them. There are, in principle, many possible ways to select a subset of peaks that cover half of the total peak probability mass; each such subset would be approximately equivalent from the perspective of maximizing information gain. In our setting, the height of each peak contains information about how likely the corresponding anchor point is to yield reward. Thus, the strategy that maximizes the probability of reward while simultaneously maximizing information gain is the one that selects the smallest number peaks that will cover half of the probability mass (this corresponds to selecting the strongest peaks). In our setting, this has the additional advantage that a smaller set of peaks corresponds to a smaller set of anchor points, which leads to simpler and more efficient navigational trajectories. Deviations from this strategy, for example induced by sampling anchors in proportion to the posterior, or by adding execution noise to these anchors, will slow the convergence of the posterior onto a successful anchor point. Given a set anchor points, the agent can execute a trajectory that them, as described above. If this trajectory leads to reward, the agent can eliminate the remaining anchors; alternatively, if the trajectory does not lead to reward, the agent can eliminate the current set of anchors. Both of these operations will happen naturally via the Bayesian update described above. In either case, the agent will be left with approximately *N/*2 peaks in its posterior, from which it can again select half of these peaks to sample on the next trial. This process will quickly narrow down the successful anchor point by eliminating anchor points that are inconsistent with reward.

### Planner

The sampling process described in the previous section can be used to determine a set of anchors that form the basis of a trajectory. We assume that this set of anchors is relayed to a planning module that uses these anchors to plan and execute a trajectory. The planner must decide the appropriate ordering of the anchors, and the precision with which the planned trajectory will intercept the anchors.

#### Ordering of anchors

We assume that the agent tries to minimize energy expenditure by planning a path that traverses the set of anchor points as efficiently as possible. To that end, we solve a simplified version of the traveling salesman problem to determine the optimal ordering of anchors. For up to five anchors, the total number of permutations is small and can be explored exhaustively; in this case, we enumerate all possible permutations, and we find the ordering that minimizes the path length of the trajectory between them. For more than five anchors, we break the problem into multiple sub-problems, solve each sub-problem using the same technique, and merge the solutions to form a complete trajectory. This can approximated by a much simpler algorithm in which anchors are ordered according to their orientation.

#### Precision of execution

Once we have determined the optimal ordering of anchors, we allow the agent to adjust the positioning of anchors, up to some tolerance, in order to further reduce the path length of its trajectory. We set this tolerance on an anchor-by-anchor and trial-by-trial basis, based on the width of the posterior peak from which the anchor was sampled. Thus, anchors that are sampled from broad peaks will have a higher tolerance—and thus can be shifted over larger distances—compared to anchors sampled from narrow peaks. This captures the idea that strong peaks in the posterior (which represent regions of greater certainty) will tether the behavior more strongly than weak peaks (which represent regions of greater uncertainty).

### Context switches: resetting, caching, and sampling from prior beliefs

#### Resetting the prior following a surprising observation

The agent’s posterior belief, when combined with information about the outcome of an executed trajectory, gives an expectation of that outcome. This can be used to evaluate the agent’s surprise about a given outcome, which can in turn be used to quickly detect changes in the environment that are inconsistent with the agent’s current belief.

Given an executed trajectory 𝒯_*n*_(*t*; {(*r*_*i*_, *θ*_*i*_)}) = {(*r*_*t*_, *θ*_*t*_)}_*n*_, and an observed outcome *o*_*n*_, Eq. 23 specifies how to update the posterior belief. The precise update involves normalizing by the sum of the distributions on the right hand side of the equation; this normalization constant specifies the probability of the current outcome given past and present trajectories, and given past outcomes (i.e., given the prior belief and the just-executed trajectory):

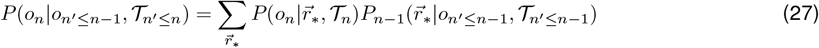

This probability can then be used to compute the surprise of the current outcome, given the prior belief and the executed trajectory:

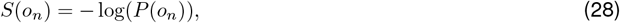

where we have dropped the conditional dependencies for ease of notation.

This signal will decrease over time as learning converges (i.e., as the agent’s belief localizes in a manner that generates trajectories that are consistent with reward). Upon a target switch, what had previously been rewarding trajectories will no longer intercept the target, and thus will no longer generate reward; this results in a large surprise signal, because the outcome is inconsistent with the current belief. The agent can use this signal to detect a switch the environment and respond accordingly. The simplest response is to reset the posterior belief to a uniform prior, which significantly speeds the subsequent learning process.

#### Storing the existing posterior in a memory cache

In many settings, it might also be advantageous to store, or “cache”, the current posterior before reverting to a uniform belief. This has the advantage of preserving the memory of a useful belief to be used again in the future.

To account for this possibility, we consider a scenario in which the agent maintains a memory of past contexts *C*. Each context corresponds to a stored *anchor belief* (i.e., a belief about the successful anchor). The agent then maintains a *context belief P* (*C*) about the current context, which it uses to select an appropriate anchor belief to guide a trajectory on a given trial. The agent can then use the executed trajectory, together with its outcome, to update both its anchor belief (as given in Equation 23 above), and to update its context belief:

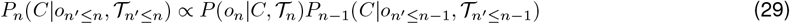

where *P* (*o*_*n*_ |*C*, 𝒯_*n*_) is the probability of receiving the outcome *o*_*n*_ given the executed trajectory 𝒯_*n*_ and the prior belief obtained from a given context *C*. In other words, this is the outcome probability given in Equation 27, where the context *C* serves as a proxy for the full anchor belief that characterizes the context. We will refer to this as the “context posterior”, to distinguish it from the posterior belief about anchor points (i.e., the “anchor posterior”).

The set of contexts is specified by the set of previously-cached anchor beliefs, together with the current anchor belief; the context posterior in Equation 29 is thus defined over this set. We assume that the agent can use either the previously-cached beliefs or the current current working belief to guide its trajectory on any given trial, but we assume that the agent only updates its current working belief based on the outcome of this trajectory. There are many possible ways to sample from the cache on a given trial, but we consider two possibilities: the agent samples the most probable context, or samples in proportion to the posterior probability of each context. The agent uses the corresponding anchor belief to sample anchor points for a given trajectory.

Because the agent can cache its current anchor belief if incoming observations are sufficiently surprising, the space of available contexts can grow over time. When a new context is created, the context posterior will grow in size and must be renormalized. Upon creating a new context, we assume that the existing context posterior is renormalized to sum to 0.5; the remaining probability of 0.5 is assigned to the newly-created context. This has the effect of prioritizing the new context relative to past contexts; the larger the existing cache, the bigger the relative difference between the probability of the new context and the next most probable context.

### Obstacle avoidance

The algorithm detailed above seeks to find a single anchor that will reliably generate a rewarding trajectory. In complex environments—such as those with obstacles or more complex geometries—such single-anchor trajectories may not be flexible enough to reliably intercept the target. To overcome this challenge, we allow the agent to augment its trajectory with additional anchor points. With this addition, we can view learning as the processing of trimming and expanding a set of candidate anchor points to stabilize rewarding trajectories.

To guide the process of augmenting trajectories, we allow the agent to track the path-integrated error between its planned trajectory, and the trajectory that it can execute in the environment. In an open arena where the agent knows the arena bounds, the planned and executed trajectories are identical. However, if an obstacle is introduced into the arena, the agent may have to divert a planned trajectory around the obstacle, leading to a difference between the planned and executed trajectory. We denote this difference as an error 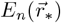, defined over the same space as the anchor belief. This error can be computed using the likelihood function given in Eq. 24:

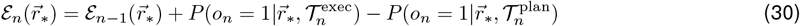

where 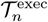 and 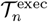 are the planned and executed trajectories, respectively. This object takes on positive values for vectors 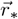 at which the executed trajectory deviated from the planned trajectory; as such, these positive regions specify candidate anchors that are likely to be successful in the future (where here, “successful” corresponds to avoiding obstacles, rather than collecting reward). Similarly, the negative regions of this object correspond to vectors 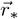 at which the planned trajectory deviated from the executed, and are thus to be avoided in the future.

We allow the agent to use this error to augment its trajectory with additional anchor points. To determine whether or not to augment a planned trajectory, the agent can evaluate whether the trajectory is likely to generate high errors. There are many ways to specify this; we choose a heuristic approach, and allow the agent to augment a planned trajectory if the following condition is met:

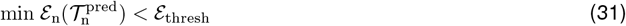

where we use the minimum of ℰ because negative error values correspond to regions where previously-planned trajectories were not successful, and thus should be avoided.

If this condition is met, we allow the agent to augment its trajectory with anchors sampled from the peaks of ℰ, corresponding to regions where previously-executed trajectories were successfuly diverted around obstacles.

### Pseudocode

Below, we detail the algorithmic steps described above.

#### Algorithm 1

Learn to plan and execute a trajectory that intercepts a hidden target

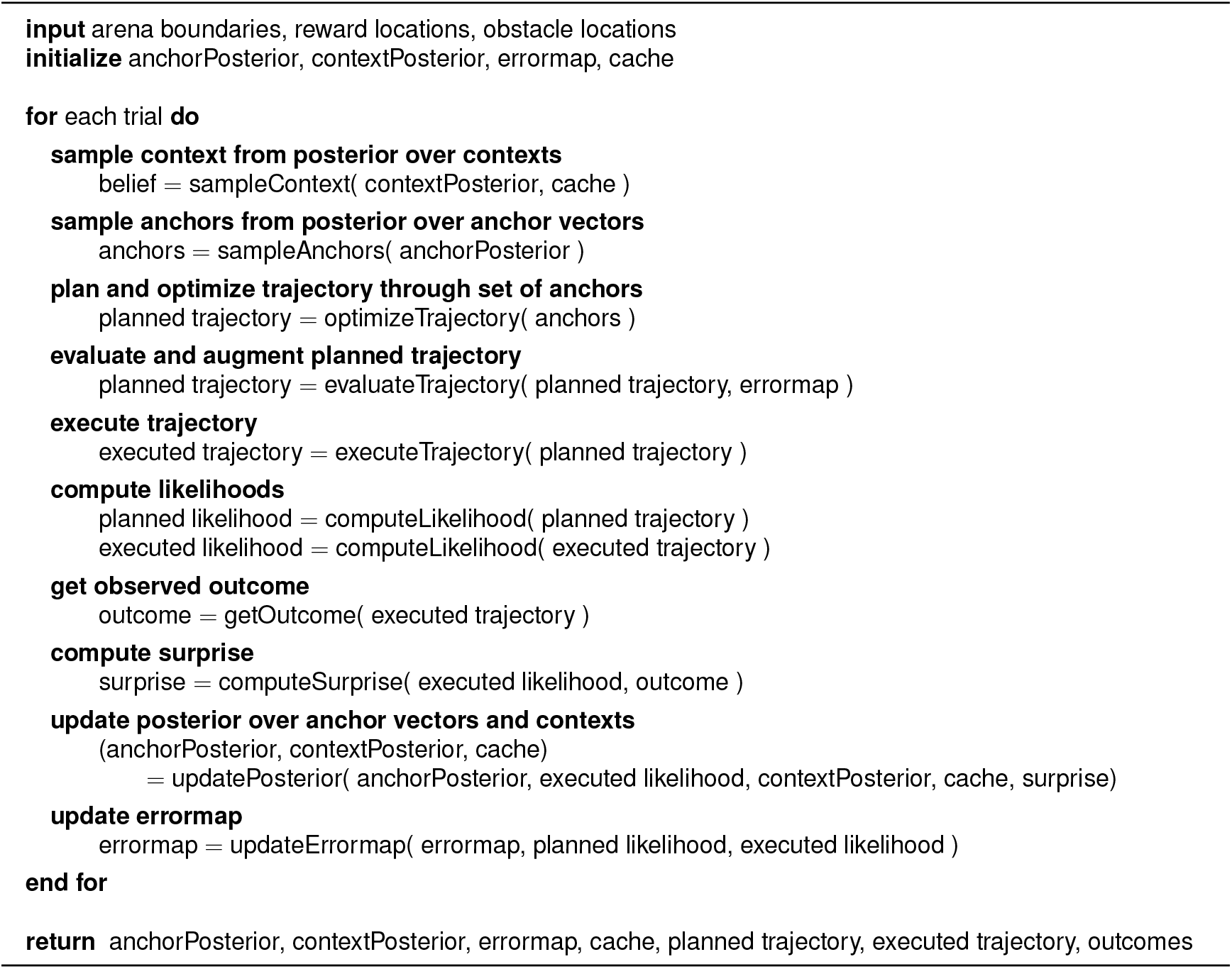

#### Algorithm 2

Sample context from posterior over contexts

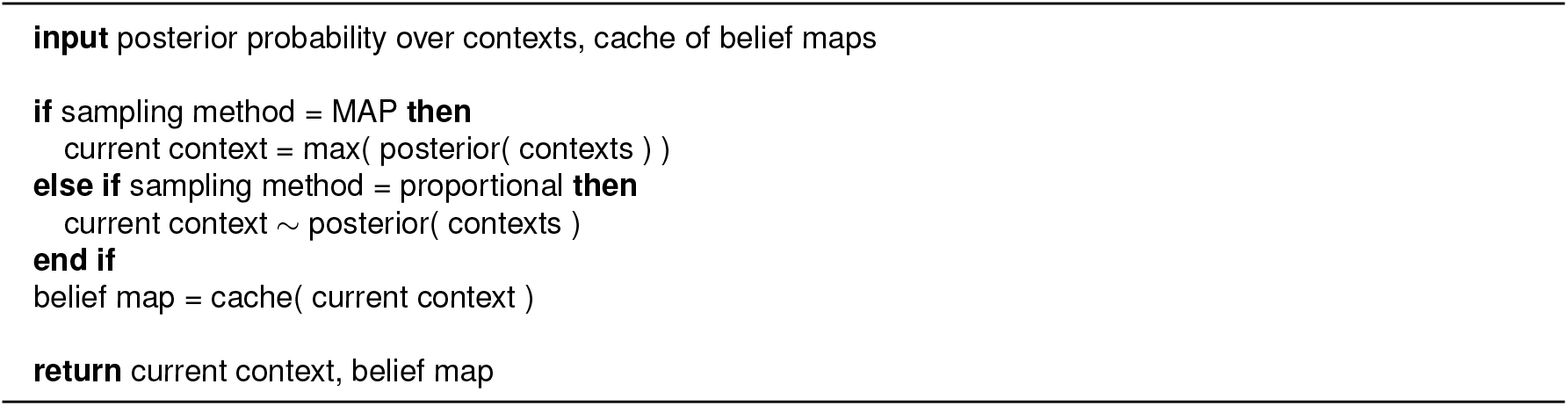

#### Algorithm 3

Sample anchors from posterior over anchors vectors

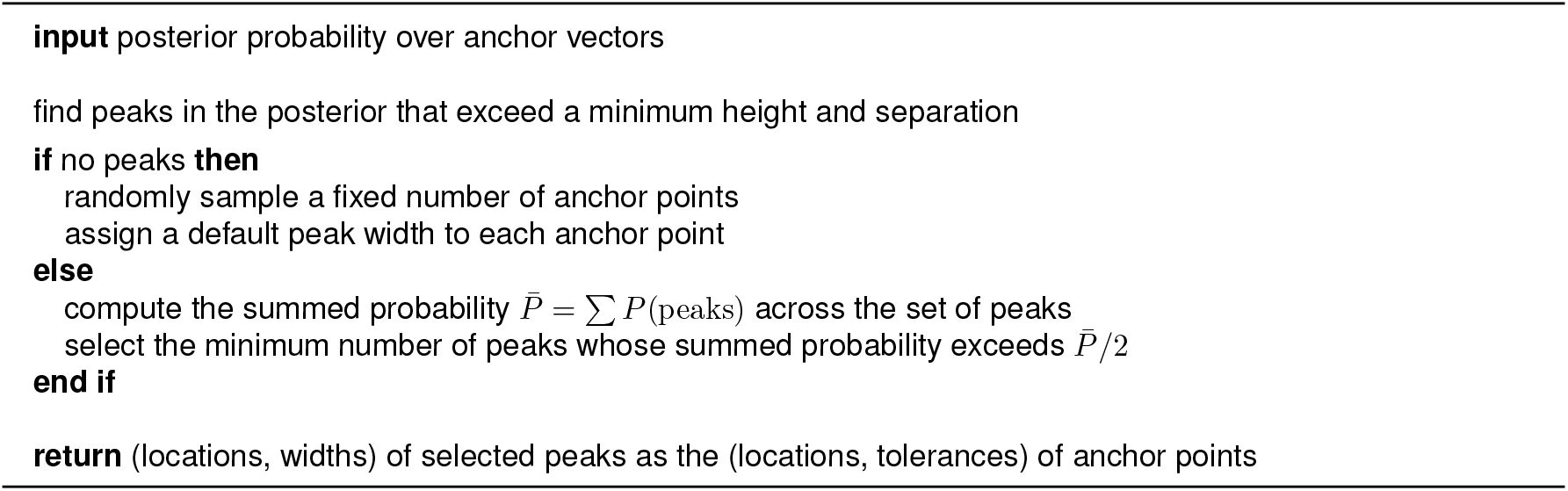

#### Algorithm 4

Optimize trajectory through set of anchors

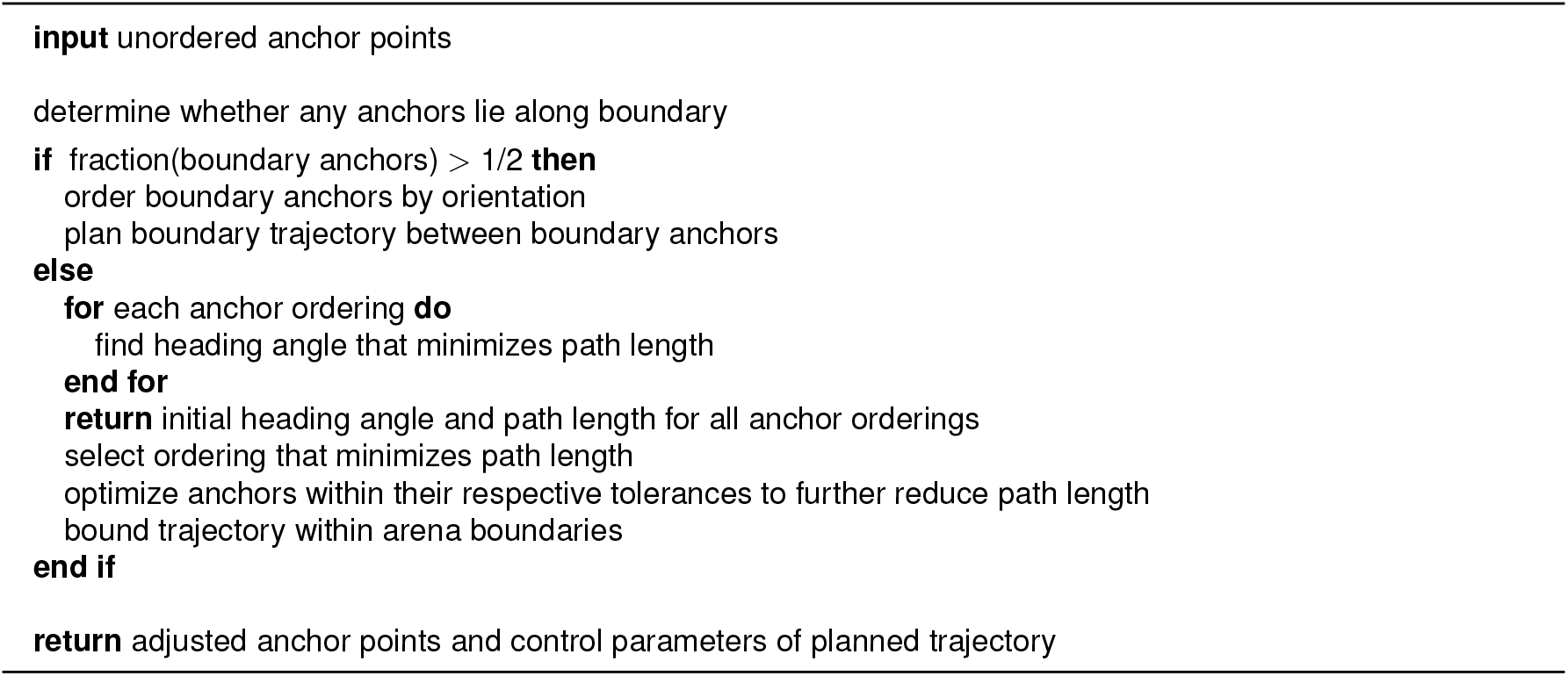

#### Algorithm 5

Plan trajectory through set of anchors

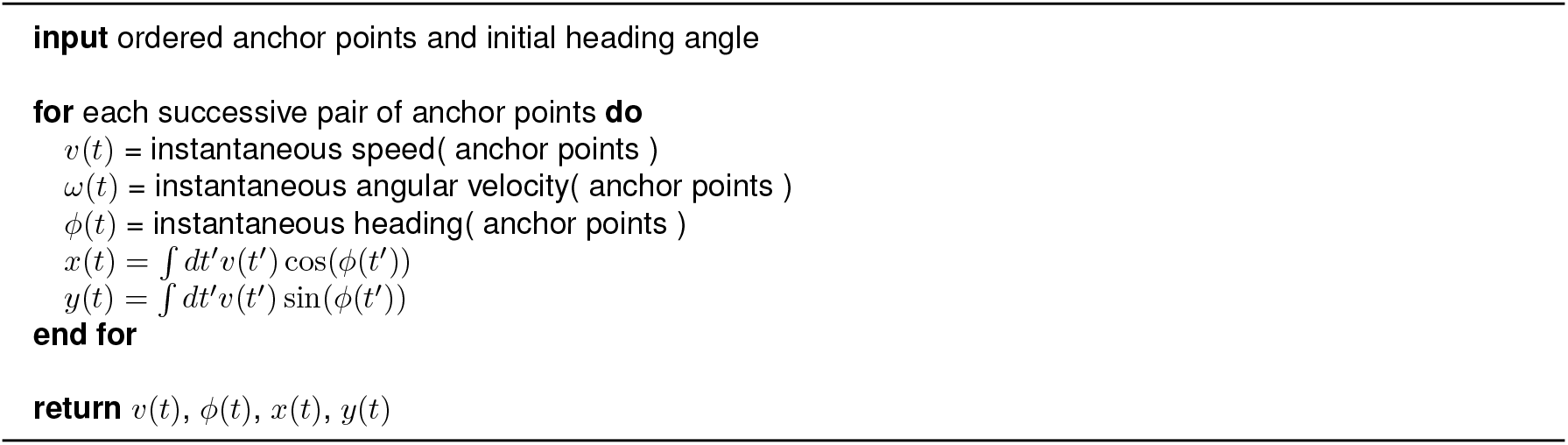

#### Algorithm 6

Evaluate planned trajectory

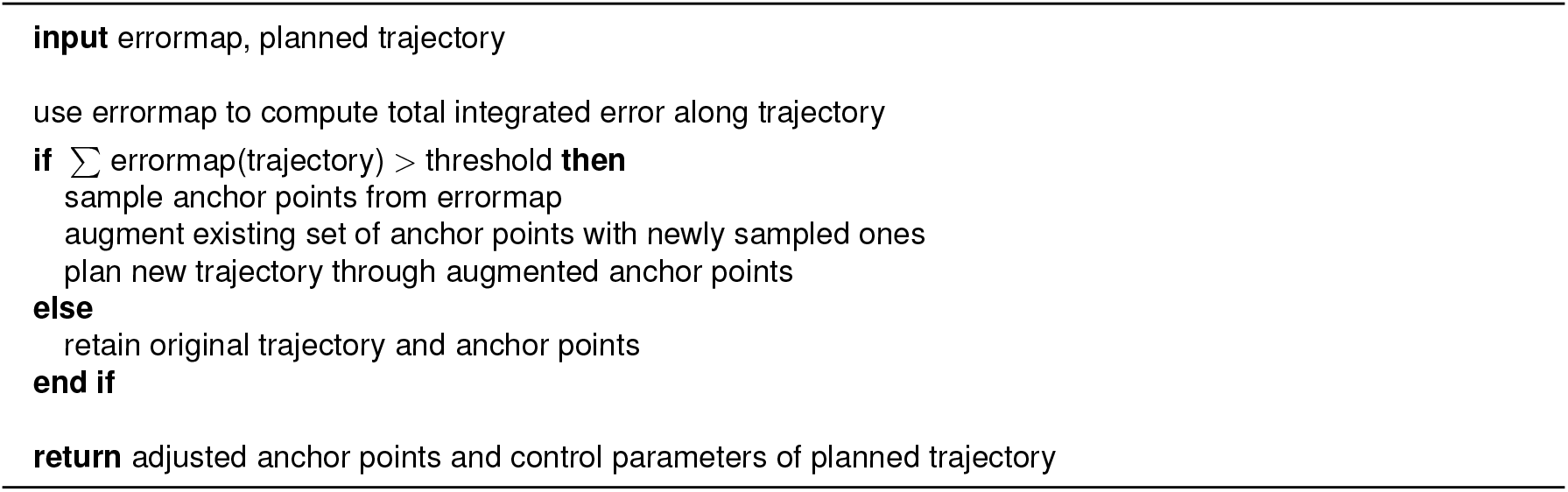

#### Algorithm 7

Execute trajectory

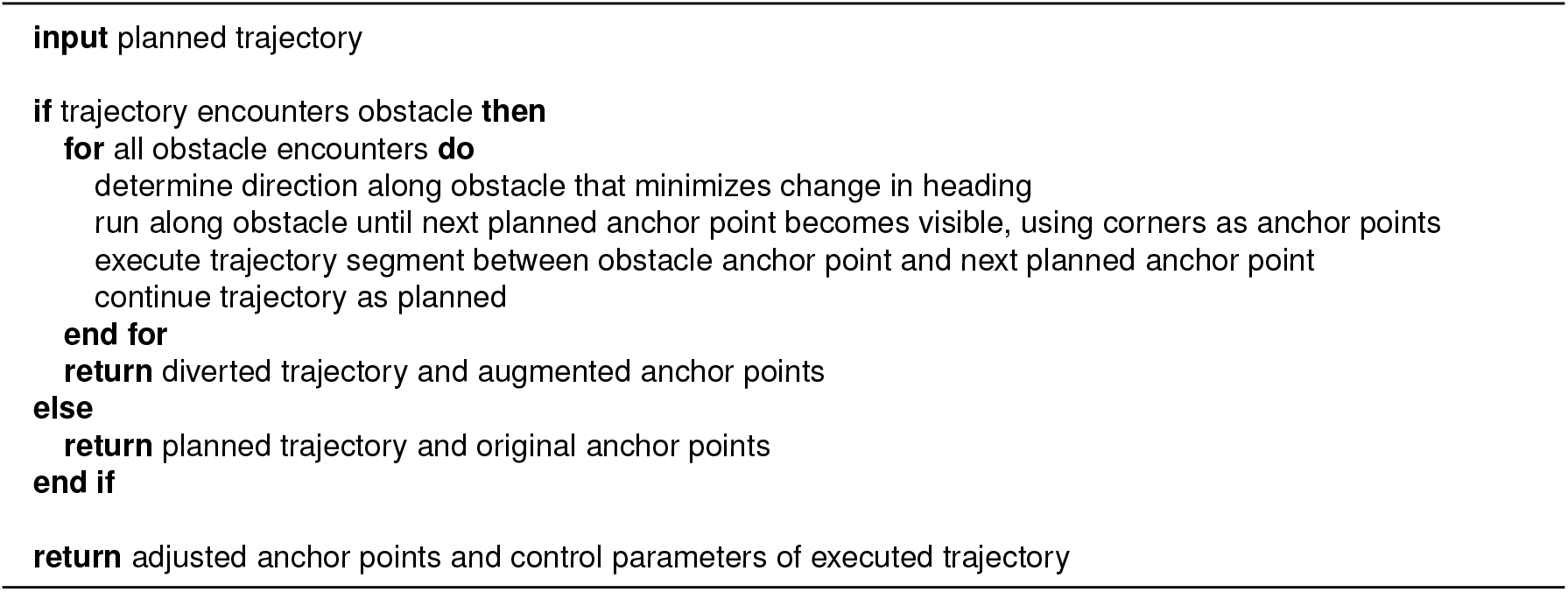

#### Algorithm 8

Compute likelihood

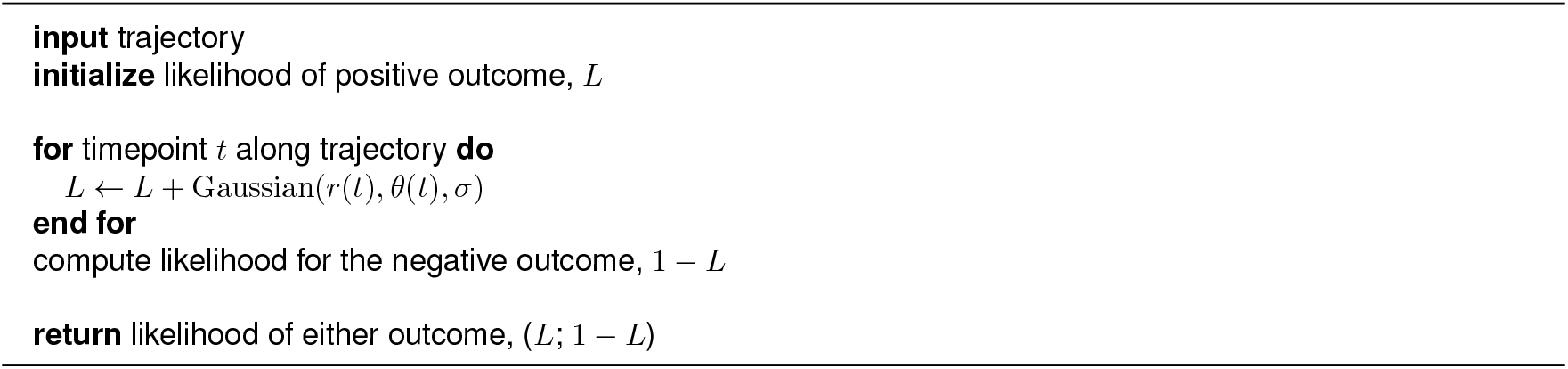

#### Algorithm 9

Get outcome

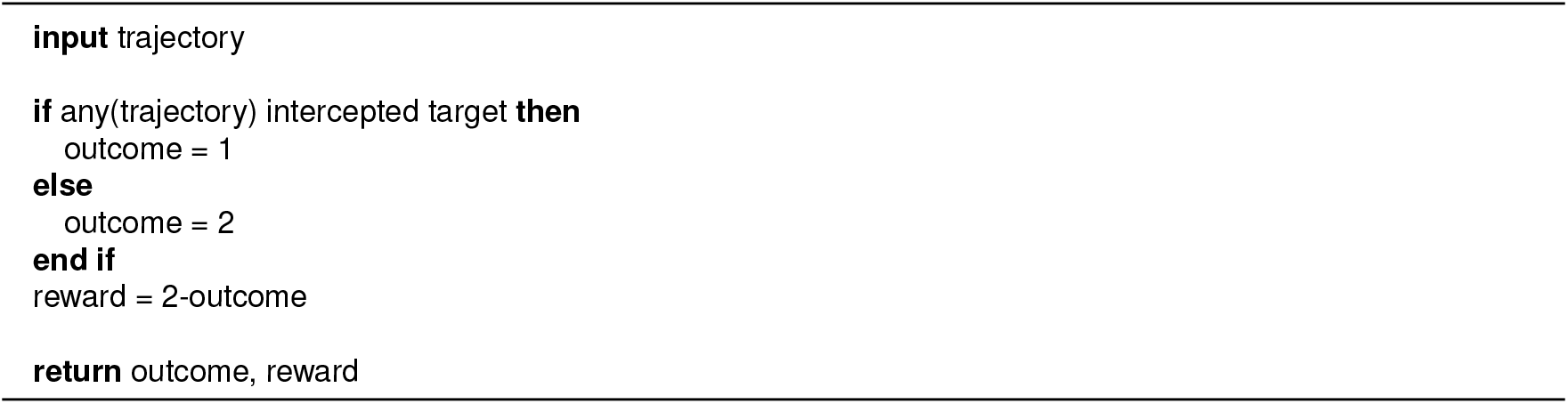

#### Algorithm 10

Compute surprise

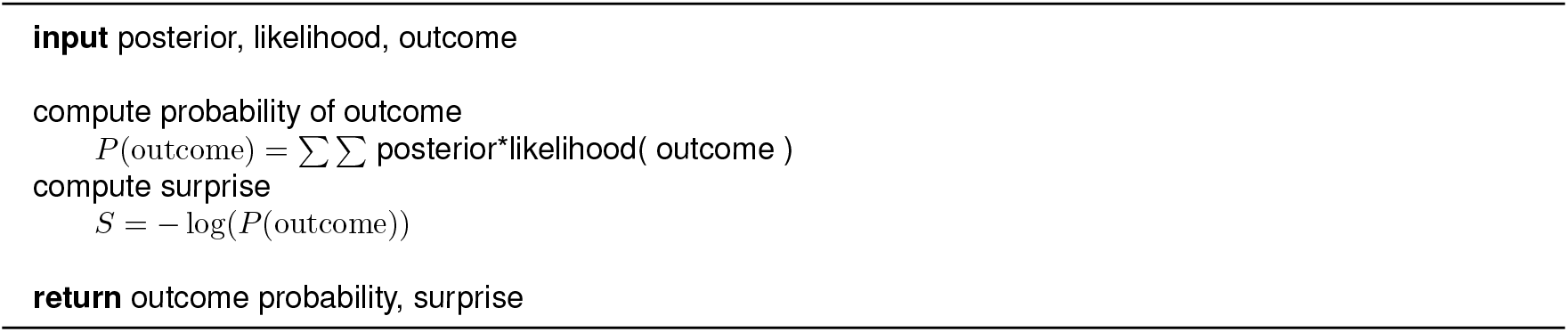

#### Algorithm 11

Update posteriors over anchor vectors and over context

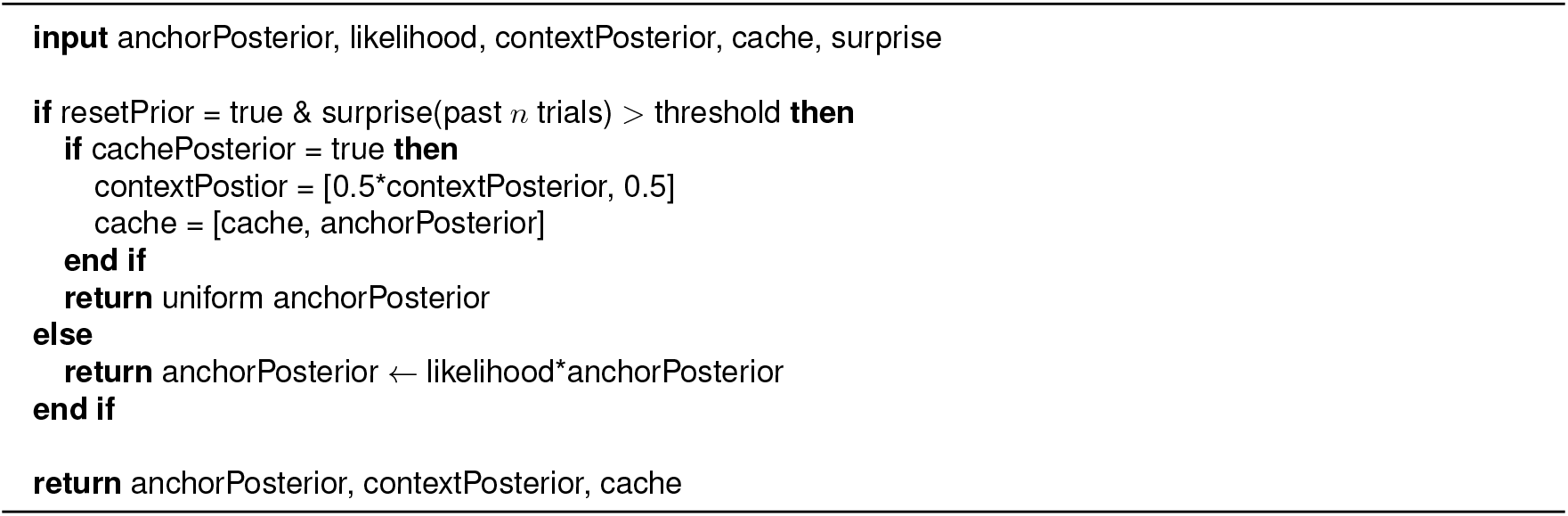

#### Algorithm 12

Update errormap

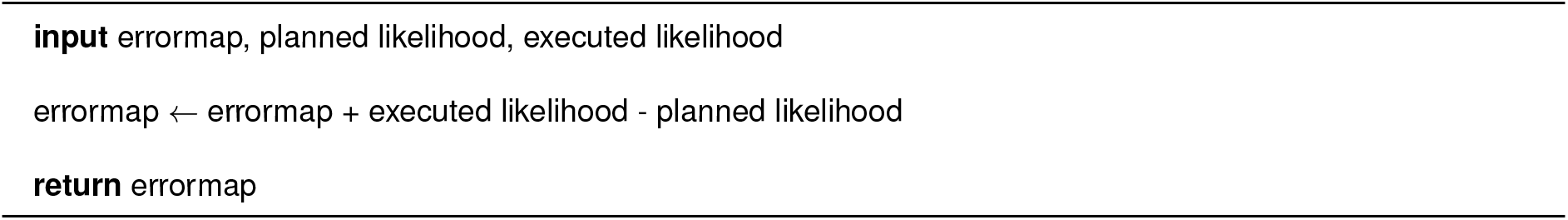

